# When do individuals maximize their inclusive fitness?

**DOI:** 10.1101/624775

**Authors:** Laurent Lehmann, François Rousset

## Abstract

Adaptation is often described in behavioral ecology as individuals maximizing their inclusive fitness. Under what conditions does this hold and how does this relate to the gene-centered perspective of adaptation? We unify and extend the literature on these questions to class-structured populations. We demonstrate that the maximization (in the best-response sense) of class-specific inclusive fitness obtains in uninvadable population states (meaning that all deviating mutant go extinct). This defines a genuine actor-centered perspective on adaptation. But this inclusive fitness is assigned to all bearers of a mutant allele in a given class and depends on distributions of demographic and genetic contexts. These distributions, in turn, usually depend on events in previous generations and are thus not under individual control. This prevents, in general, from envisioning individuals themselves as autonomous fitness-maximizers, each with its own inclusive fitness. For weak selection, however, the dependence on earlier events can be neglected. We then show that each individual in each class appears to maximize its own inclusive fitness when all other individuals exhibit fitness-maximizing behavior. This defines a genuine individual-centered perspective of adaptation and justifies formally, as a first-order approximation, the long-heralded view of individuals appearing to maximize their own inclusive fitness.

## Introduction

One striking hallmark of living systems is their functional organization. From molecular, cellular, and physiological structures within individuals to behavioral interactions between them, organisms in nature display a purposefulness in form and a goal-directedness in action that has been marveled at by generations of biologists (Darwin, 1859; Fisher, 1930; Williams, 1966; Dawkins, 1996; Grafen, 2007). This outward functionality is so unequivocal that humanity has attributed purpose to animals and plants since the mists of time.

Can this purposefulness be characterized? It is well-understood that the functionality of organisms is born out of natural selection. This causes organisms to become adapted to their biotic and abiotic environments over evolutionary time. Over short time scales, mutations are limited and allele-frequency changes, resulting from differences in organismic forms and behaviors, involve selection among a limited number of alternative variants present in the population. Since to each trait combination of an organism there is an associated reproduction and survival schedule, the process of genetic adaptation is often depicted as the maximization of individual fitness. Survival and reproduction, however, also depend on the environment in which individuals reside, and in particular on the traits of conspecifics. An organism’s environment thus varies in response to changes in trait composition in the population induced by natural selection. This prevents a net increase in individual fitness over evolutionary time, even supposing that at all times alleles increasing survival and reproduction are favored by evolution. Indeed, the goalposts of the survival and reproductive games of life are shifting as evolution proceeds. And even with fixed goalposts, increase in survival and reproduction may be prevented by multilocus effects in the presence of recombination, as highlighted in some classical criticisms of fitness maximization (e.g., Moran, 1964; Ewens, 2004; Bürger, 2000; Ewens, 2011).

Over long time scales, an organism can be regarded as adapted to a particular environment if no alternative trait combination or behavioral schedule can be produced by mutation, which would result in further allele frequency change (Fisher, 1930; Williams, 1966; Grafen, 1988; Reeve and Sherman, 1993; Dawkins, 1996). In this long-term perspective, the maximization of the geometric growth ratio of a mutant allele when rare–referred to here as *invasion fitness*–in a large population where individuals express a resident allele, provides a condition of uninvadability of mutant traits (all deviating mutants from some feasible set of traits go extinct). Uninvadability is a defining property of an evolutionary stable population state in which the resident trait combination is a best-response to any mutant deviation (Eshel, 1983; Metz et al., 1992; Ferrière and Gatto, 1995; Eshel et al., 1998; Metz, 2011). It is in terms of this notion of best-response that maximization of (invasion) fitness can actually be conceived in the long-term evolutionary perspective and this holds regardless of the underlying genetic details (Eshel and Feldman, 1984; Liberman, 1988; Eshel, 1996; Hammerstein, 1996; Weissing, 1996; Eshel et al., 1998).

Invasion fitness is the per capita number of mutant copies produced by the whole mutant lineage descending from an initial mutation over a life-cycle iteration, when the mutant reproductive process has reached stationarity in a resident population. This shows that invasion fitness is a property of a collection of interacting individuals, and gives no reason to say that in an uninvadable population state the fitness of any of these individuals is maximized (in the best-response sense). It has even been argued that any focus on individual survival and reproduction to understand adaptation is misleading and should be abandoned altogether (Dawkins, 1978). The gene-centered perspective of adaptation (Hamilton, 1963, 1996; Dawkins, 1976, 1982; Haig, 1997b, 2012) has in fact distanced itself from ideas of maximization of individual survival and reproduction long ago, and focuses instead on the differential transmission of alleles to understand adaptation.

In spite of the logical primacy of the gene-centered perspective, trying to interpret natural selection in individuals terms appears necessary for anyone observing individuals rather than genes. Hamilton (1964) attempted to draw a bridge between the gene and the individual-centered perspective of adaptation by defining inclusive fitness, a quantity that is assigned to a representative carrier of an allele, so that natural selection proceeds as if this quantity is maximized. While this inclusive fitness is not a distinct property of each individual in a population, individual inclusive fitness maximization has nevertheless been argued to have at least some heuristic value (e.g., Maynard Smith, 1982; Dawkins, 1978; Grafen, 1984, 2007; West and Gardner, 2013), and is a working assumption in behavioral ecology (McNamara et al., 2001; Alcock, 2005) and evolutionary psychology (Alexander, 1990; Buss, 2005). It is also a perspective often endorsed in social evolution theories (Bourke, 2011; West and Gardner, 2013). For example, one may say that sterile workers maximize their inclusive fitness by helping a colony queen to raise offspring. Here, it is acknowledged that workers, being sterile, do not maximize their individual fitness, but rather the survival and expected reproduction of related individuals.

Despite the attractiveness of the individual-centered perspective of adaptation, there has been few formal models supporting it and/or delineating the conditions under which individuals can be regarded as autonomous agents maximizing their own objective function (i.e., maximizing their own maximand). For instance, Grafen (2006a) considers that individuals maximize an inclusive fitness, which, formally, does not depend on the behavior of other conspecifics and thus appears on our reading not to cover social interactions in any broad sense. It has also been shown that, in age-structured population without social interactions and in group-structured populations with social interactions, individuals appear to maximize a weighted average individual fitness (respectively Grafen, 2015 and Lehmann et al., 2015), which is distinct from inclusive fitness. More generally, connections between individual fitness, inclusive fitness, and individual maximization behavior have been discussed in the literature on kin selection, evolutionary stable traits, and adaptive dynamics (e.g., Hines and Maynard Smith, 1978; Michod, 1982; Maynard Smith, 1982; Eshel, 1991; Mesterton-Gibbons, 1996; Eshel et al., 1998; Day and Taylor, 1996; Frank, 1998; Day and Taylor, 1998; Rousset, 2004; Lehmann and Rousset, 2014a; Akçay and Van Cleve, 2016; Okasha and Martens, 2016; Eshel, 2019). But these discussions often do not emphasize enough the distinction between the gene-centered and the individual-centered perspective of adaptation, and generally do not cover the case of class-structured populations (e.g., queen and worker, male and female, young and old individuals).

Our goal in this paper is to formalize and push forward, as far as is consistent with the gene-centered perspective, the individual-centered approach according to which individuals may maximize their own inclusive fitness in an uninvadable population state. To that aim, we use the common framework of evolutionary invasion analysis and proceed as follows in four steps. We start by presenting a model of evolution in a diploid group-structured population with limited dispersal and class-structure within groups. In this model, we then consider three perspectives on adaptation, formalized by three fitness measures, all of which are maximized (in the best-response sense) in an uninvadable population state. First, we consider a “recipient-centered perspective”, which is formalized as a weighted mean of the fitness of individuals who bear an allele. Second, we consider an “actor-centered perspective”, which is formalized as an exact class-specific version of inclusive fitness. This maximand is assigned to all bearers of an allele in a given class. Finally, we consider a “rational actor-centered perspective”. This is the only perspective that fully captures the idea that individuals maximize *their own* inclusive fitness, and is formalized as a class-specific inclusive fitness that can be assigned to individuals without knowing their genotype.

By formalizing these three perspectives on adaptation and discussing similarities and differences thereof, we unify and extend previous results on the relationship between maximization behavior and the concept of adaptation *sensu* Reeve and Sherman (1993, p. 9), i.e., “A phenotypic variant that results in the highest fitness among a specified set of variants in a given environment”. This allows us to provide a full connection between evolutionary invasion analysis, the different perspectives on inclusive fitness theory, and game-theoretic approaches. Readers who find the technical details in the following text challenging may nevertheless have a look at the expression for inclusive fitness (eq. 4) and read the Discussion. Therein, results are summarized with a number of take-home messages about the interpretation of adaptation in terms of fitness maximization. Finally, readers interested in how selection on traits can be decomposed into direct effect on actors and indirect effects on recipients, but not on whether individuals do really maximize their own inclusive fitness, should read the section “The actor-centered perspective of adaptation”, but can skip the more complicated section “The rational actor-centered perspective of adaptation”.

## The model

### Assumptions

In order to formalize and compare the different perspectives on adaptation in a simple way but retain key biological population structural effects, we endorse two sets of well-studied assumptions.

#### Demographic assumptions

First, we assume that evolution occurs in a population structured into an infinite number of groups (or demes or patches), each with identical environmental conditions, and connected to each other by random and uniformly distributed, but possibly limited, dispersal (i.e., the canonical demographic island model of Wright, 1931). Demographic time is discrete and during each demographic time period, reproduction, survival, and dispersal events occur in each group with exactly *n* individuals being censused at the end of a time period (after all relevant density-dependent events occurred). Each of the *n* individuals in a group belongs to a class (e.g., queen and worker, male or female, young or old) and *n*_c_ denotes the number of classes, which is assumed fixed and finite. The number of individuals in each class can differ within a group (but this difference is the same for all groups).

Each individual in each group can express a class-specific trait that affects its own survival, reproduction, and dispersal and possibly those of group neighbors. Let us focus on a focal group where individuals are labelled from 1 to *n* and let *x*_*i*_ denote the trait expressed by focal individual *i* from that group. When this individual is of class *a*, its trait is taken from the set 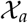 of feasible traits available to an individual of class *a* 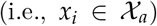. We collect the class-specific traits expressed by all of the *n* − 1 neighbors of *i* into the vector ***x***_−*i*_ of group-neighbor traits. We then let 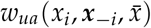 denote the expected number of surviving class-*u* offspring *per haplogenome* produced over a demographic time period by a class-*a* individual *i* with trait 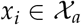 when group neighbors have trait profile ***x***_−*i*_ in a population where the average individual trait profile across classes is 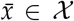 (here 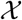 denotes the set of feasible traits across all classes; see Supplement A, eq. A.3, for a formal definition of fitness and Table 1 for a summary of notation).

We refer to *w*_*ua*_ as the *individual fitness* function as it determines the number of successful gametes per haploid set of an individual (Grafen, 1985). Individual fitness thus gives the average number of replicate gene copies produced by an individual per homologous gene and so the unit of measurement is here gene copy number. In writing individual fitness as 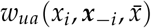 we made, for simplicity of analysis and presentation, a number of assumptions on fitness. First, individual fitness depends here on the focal’s own phenotype only through the trait of the class in which the focal resides. This thus excludes interdependence among own traits throughout the lifespan of the focal when it can change class (such as in an age-structured population when traits affect body size and thus may have effects on different age classes). Second, the focal’s individual fitness depends on the trait of each neighbor only through the neighbor’s class-specific trait. This thus also excludes interdependence among traits throughout the lifespan of neighbors that can change class (such as in an age-structured population when traits affect body size and transfer of resources among individuals depends on body size). Finally, the effects of individuals from different groups on a focal individual’s fitness is mean-field; that is, it depends only on the population average trait. While making these assumptions on fitness do not allow us to cover all biological scenarios of interest (see section “Scope of our results” for a discussion of the restrictive assumptions of our model and how to relax them), our model does cover the evolution of traits of arbitrary complexity in disjoint classes (i.e. when the class of an individual is fixed at birth), like workers and queens or males and females. As such, our model applies in broad generality to the paradigmatic biological scenarios for which inclusive fitness theory has been initially developed.

#### Evolutionary assumptions

No assumptions so far have been made on the genetic composition of the population and each individual may express a different trait and thus be phenotypically distinct from any other individual. In order to understand which traits are favored over the long term by evolution in this population, we now turn to our second set of assumptions. We place ourselves in the framework of an evolutionary invasion analysis (“ESS approach”, e.g., Eshel and Feldman, 1984; Tuljapurkar, 1989; Parker and Maynard Smith, 1990; Metz et al., 1992; Charlesworth, 1994; Ferrière and Gatto, 1995; Eshel, 1996; Caswell, 2000; Otto and Day, 2007; Metz, 2011). Accordingly, we consider a population that is monomorphic for some resident trait and aim at characterizing the conditions according to which a mutant allele changing trait expression is unable to invade the population. For this, we let resident individuals (necessarily homozygotes if diploid) have the vector 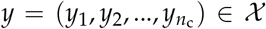 of traits, one for each class of individuals, where *y*_*a*_ is the trait of a (homozygote) individual of class *a*. Let a heterozygote mutant individual have trait vector 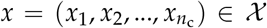. We assume that heterozygote traits are convex combinations (“weighted averages”) of homozygote traits, which means that we rule out over-, under-, and strict dominance, but otherwise allow for arbitrary gene action. This allows us to write the trait *z*_*a*_(*x*_*a*_, *y*_*a*_) of mutant homozygotes of class *a* as a function of the traits *x*_*a*_ of heterozygote and *y*_*a*_ of resident homozygotes of that class (Supplement eqs. A.13–A.14 for a formal definition). This also covers the haploid case where we simply assign trait *x* to mutant and trait *y* to residents. So, for both haploid and diploid populations, traits are fully specified by *x* and *y*.

## Characterizing adaptation by way of uninvadability

With the assumptions of the last section, the fate (extinction or spread) of a mutant allele with heterozygote trait *x* introduced into a monomorphic resident class-structured population with homozygote trait *y* can be determined by the *invasion fitness W*(*x*, *y*). This is the geometric growth ratio of the mutant allele when rare. Henceforth, the mutant allele cannot invade the population when

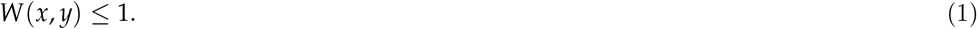

Suppose now that a given resident trait, say 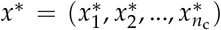, is uninvadable. Namely, it is resistant to invasion by any alternative trait from the set of all possible traits 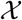 (i.e., eq. 1 holds for any mutant given the resident *x*^∗^). This trait *x*^∗^ must then be a best response to itself, meaning that if we vary invasion fitness *W*(*x*, *x*^∗^) by varying *x*, the uninvadable trait *x*^∗^ must maximize invasion fitness with respect to 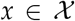. This means that *x*^∗^ results in the highest invasion fitness among all alternatives traits given in the set 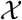 of feasible traits, for the resident population at the uninvadable state. Hence, *x*^∗^ qualifies as an adaptation in the sense of Reeve and Sherman (1993, p. 9), i.e., “A phenotypic variant that results in the highest fitness among a specified set of variants in a given environment” with “fitness” being invasion fitness and “a specified set of variants” being 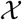. This definition of adaptation is useful for two reasons. First, it has a direct formal encapsulation as uninvadability. Second, it captures the apparent purposefulness of traits as an outcome of cumulative natural selection, which has forcefully been argued as being the defining feature of adaptation (Fisher, 1930; Williams, 1966; Grafen, 1988; Dawkins, 1996).

The above characterization of adaptation is gene-centered, since it is obtained in terms of the fitness of mutant alleles. In order to assess whether individuals maximize their inclusive fitness, however, we must characterize adaptation in terms of individual-centered concepts. For that purpose, we contrast three individual-centered perspectives on adaptation:

**Perspective (1), recipient-centered.** Here, we provide a representation of invasion fitness expressed in terms of the class-specific fitness components (the *w*_*ua*_ functions introduced above) of a typical carrier of the mutant allele. This perspective directly flows out from the evolutionary model and is recipient-centered because we consider the fitness of individuals bearing the mutant allele and how this fitness is affected by the behavior of others. Hence, behavioral effects are grouped by recipients.
**Perspective (2), actor-centered.** Behavioral effects, however, are the outcomes of actors expressing traits. We thus ask whether it is possible to obtain a class-specific representation of fitness that is maximized (in the best-response sense) at uninvadability and where behavioral effects are grouped by the actor. That is, can we define a (class-specific) inclusive fitness that is maximized for each class in an uninvadable population state?
**Perspective (3), rational actor-centered.** Here, we go one step further than the actor-centered perspective. We ask whether it is possible to obtain an actor-centered representation of fitness where the trait of each individual is different and where this fitness is still maximized at uninvadability, independently for each individual in each class. This is the (behavioral ecology) question of whether we can we look at adaptation as if each individual is an autonomous decision-maker having free choice of action and *as if* these actions are guided by a striving to maximize a measure of inclusive fitness.

Perspective (1) obtains when an allele’s fitness is viewed as a (possibly weighted) mean of the fitness of individuals who bear the allele. It is therefore a straightforward translation of the gene-centered perspective in individual-centered terms; in the context of this paper, it is hardly distinguishable from the gene-centered perspective. Perspective (2) changes the focus from the own fitness of allele bearers to their accumulated fitness effects on different recipients. This is the classic inclusive-fitness interpretation introduced by Hamilton (1964). However, in that interpretation, a single value (the class-specific inclusive fitness) is still assigned to all bearers of the allele in a given class. By contrast, only perspective (3) may capture the idea of individuals maximizing *their* inclusive fitness, and thus it is perspective (3) that seems most often invoked in behavioral ecology to understand adaptation. We now develop each perspective in turn to assess their relative levels of generality. We formally describe these perspectives and all arguments subtending our analysis in an extensive Supplement, which fully details, generalizes, and discusses more in depth technical concepts. The Supplement also contains some additional material and references as our analysis connects to many different ideas and intertwined concepts (for instance inclusive fitness itself can be formalized in different ways, see Supplement B). The main text presents the key formalization and results of our analysis.

## The recipient-centered perspective of adaptation

### Average direct fitness

In Supplement A, we provide the relationship of invasion fitness to the individual fitness components *w*_*ua*_ by suitably averaging over the distribution of class states in which a carrier of the mutant allele can reside. We denote this distribution by the vector ***ϕ***(*x*, *y*) whose *a*th entry is the probability *ϕ*_*a*_(*x*, *y*) that a random copy of the mutant allele finds itself in class *a*. Invasion fitness of the (heterozygote) mutant *x* in a resident *y* can then be written

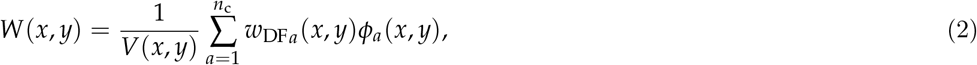

where the *average direct fitness w*_DF*a*_(*x*, *y*) of a class-*a* mutant (with “DF” standing for direct fitness) is a weighted average 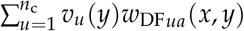, over descendants of class *u*, of the expected number *w*_DF*ua*_(*x*, *y*) of *u*-type offspring produced (per haplogenome) by an individual of class *a* bearing a copy of the mutant allele; *v*_*u*_(*y*) is the neutral reproductive value of a gene copy in class *u* (i.e., the asymptotic contribution to the gene pool of a single class-*u* gene copy in a monomorphic resident population); and *V*(*x*, *y*) is the average of the *v*_*u*_(*y*) reproductive values over the ***ϕ***(*x*, *y*) distribution, thus giving the asymptotic contribution, to the gene pool, of a randomly sampled mutant copy that is assigned the total offspring (reproductive) values of a resident gene copy. Hence, invasion fitness can be represented as the average direct fitness of a copy of the mutant allele relative to the average direct fitness this allele copy would have if it was assigned the individual fitness components of resident individuals.

We can express *w*_DF*ua*_(*x*, *y*) as an average 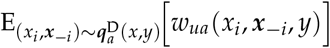 over the distribution 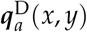 of group trait profiles (*x*_*i*_, ***x**_−i_*) experienced by a mutant gene copy [in a diploid population in particular, the trait *x*_*i*_ of the carrier of that gene copy of class *a* can be either that of a homozygote or heterozygote (*x*_*i*_ ∈ {*z*_*a*_(*x*_*a*_, *y*_*a*_), *x*_*a*_}), while the trait *x*_*j*_ of any neighbor *j* of class *s* may takes values in {*z*_*s*_(*x*_*s*_, *y*_*s*_), *x*_*s*_, *y*_*s*_}]. The distribution 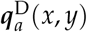 accounts for correlated phenotypic effects within groups due to identity-by-descent experienced by a carrier of class *a* of the mutant allele. Hence, the 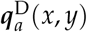 distribution captures kin selection effects experienced by a class-*a* individual, which occurs whenever the fecundity and survival of an individual is affected by the genetic trait expressed by one or more other individuals who are genetically related to the actor in a nonrandom way at the loci determining the trait (Michod, 1982, p. 40). Invasion fitness depends on the average direct fitness of each class and thus on the collection 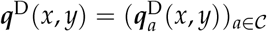 of class-specific genotype distributions. In particular,

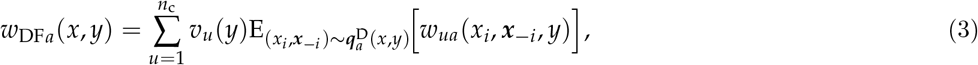

which means that the average direct fitness of a class-*a* mutant is the neutral reproductive value-weighted average of the average, over the distribution of trait profiles (*x*_*i*_, ***x***_−*i*_) experienced by a mutant gene copy, of the expected number, given the group genotypes, of *u*-type offspring produced per haplogenome by an individual of class *a* bearing a random copy of the mutant allele (see Box 1 for an example of average direct fitness).

### The importance of genetic contexts

The expression for invasion fitness *W*(*x*, *y*) (eqs. 2–3) makes explicit that invasion fitness is the average individual fitness component *w*_*ua*_ over a distributions of demographic states (captured by ***ϕ***(*x*, *y*)) and genetic states (captured by ***q***^D^(*x*, *y*)) of a copy of the mutant allele. Importantly, these two distributions depend on mutant and resident traits. As such, invasion fitness depends on the fitness of a collection of individuals taken over multiple generations and represents the average replication ability of a randomly sampled allele from the mutant lineage. This focus on gene replication epitomizes the gene-centered perspective of evolution. Accordingly, it is not the individual fitness of a single individual (or a single gene copy) in a given demographic and genetic context that matters for selection, but the average of such individual fitnesses over a distribution of contexts (Dawkins, 1978; Haig, 1997b, 2012). Natural selection on an allele thus depends not only on how it changes the immediate survival and reproduction of its carriers (changes in *w*_*ua*_), but also on changes in the probabilities of *contexts* (Kirkpatrick et al., 2002, p. 1728) in which the allele at a given locus can be found [as measured by ***ϕ***(*x*, *y*) and ***q***^D^(*x*, *y*)]. Indeed, an allele can reside, say in a worker or a queen, be inherited from a mother or a father, and is likely to be in a genome with many other loci with different allele combinations. The importance of such contexts of alleles for their evolutionary dynamics has been much emphasized in populations genetics (e.g., Altenberg and Feldman, 1987; Kirkpatrick et al., 2002; Roze, 2009). There are even mutations that spread through selection only by way of their effects on changes of the contexts in which they are found. Typical examples are modifier alleles involved in the evolution of recombination or migration, which may spread by increasing their chance of being in a genetic context with higher survival or reproduction, despite the modifier having no direct physiological effect on reproduction and/or survival in a given context (e.g., Altenberg and Feldman, 1987; Kirkpatrick et al., 2002; Roze, 2009). From now on, we refer to ***ϕ***(*x*, *y*) and ***q***^D^(*x*, *y*) as the contextual distributions.

## The actor-centered perspective of adaptation

### Inclusive fitness

What is missing to understand how selection targets traits in the representations of invasion fitness given by average direct fitness, eqs (2)–(3), is twofold. First, it is a simple and intuitive quantification of the effect of limited dispersal (and thus kin selection), which summarizes the variation on fitness introduced by the distribution over genetic contexts (the ***q***^D^(*x*, *y*) distribution). Second, it is a quantification of the contribution to fitness of trait expressions in the different classes of individuals who really contribute to allele transmission. For instance, in the social insect example described in Box 1, a male has positive individual fitness but its trait is not under selection, while a worker has zero individual fitness, but its trait is affected by selection. Then, can one identify the force of selection on the actor’s trait in a fitness measure? In other words, we aim to find an *inclusive fitness w*_IF*a*_(*x*, *y*) for a class-*a* carrier of the mutant allele in a resident *y* population, which, by varying the class-specific trait 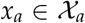 maximizes the function *w*_IF*a*_ with respect to that trait and for any actor class *a* in an uninvadable population state *x*^∗^. Hence, such maximization means that 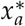 results in the highest class-*a* inclusive fitness among all traits values in 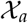 in an uninvadable population at *x*^∗^.

In Supplement B, we show that

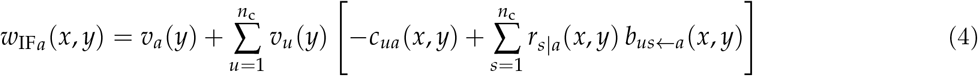

satisfies this maximization problem. Here, extending an established population genetics terminology, *−c*_*ua*_(*x*, *y*) is the *average effect* on the number of class-*u* mutant gene copies produced by a single class-*a* individual when expressing a copy of the mutant instead of the resident allele (originally, the average effect of an allele substitution on a quantitative trait; e.g., Fisher, 1941, Ewens, 2004, p. 63), and *b*_*us←a*_(*x*, *y*) is the average effect on the expected number of class-*u* offspring produced (per haplogenome) by all class-*s* neighbors in a group, and stemming from a single class-*a* gene copy switching to expressing a copy of the mutant instead of the resident allele. These costs and benefits hold regardless of the number of group partners, and have been reached by using a multiplayer and class-specific generalization of the two-predictor regression of individual fitness used in the exact version of kin selection theory (Queller, 1992; Frank, 1997; Gardner et al., 2011; Rousset, 2015; see Supplement B for further considerations on regressions and inclusive fitness). As such, *−c*_*ua*_(*x*, *y*) and *b*_*us←a*_(*x*, *y*) describe additive effects on fitness resulting from two distinct gene substitutions, as if each was independently brought up by mutation. Each such effect accounts for changes in individual fitness when everything else is held constant, in particular holding constant the average effects of interactions between individual traits (although those will be modified by an allelic substitution).

Finally, inclusive fitness (eq. 4) depends on

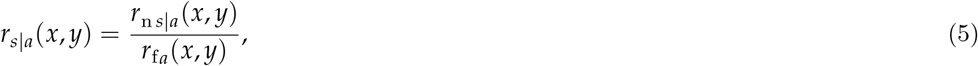

which is the relatedness between a class-*a* actor and a class-*s* recipient. Here, *r*_f*a*_(*x*, *y*) is the probability that, conditional on an haplogenome in a focal individual of class *a* carrying the mutant allele (hence the subscript “f”), a randomly sampled homologous gene in that individual is mutant, and *r*_n*s|a*_(*x*, *y*) is the probability that, conditional on an individual of class *a* carrying the mutant allele, a randomly sampled homologous gene in a (non-self) neighbor of class *s* is a mutant allele (hence the subscript “n”). Hence, relatedness *r_s|a_*(*x*, *y*) can be interpreted as the ratio of the probability of indirect transmission by a class-*s* individual of a mutant allele taken in a class-*a* individual to the probability that the individual transmits itself this allele to the next generation. In the absence of selection, this is equivalent to the standard ratio of probabilities of identity-by-descent (Hamilton, 1970, p. 1219, Lehmann and Rousset, 2014b, eq. A.5). Relatedness is expressed in terms of the class-specific mutant copy number distribution (the 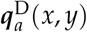 distribution) and as such summarizes the statistical effects of limited dispersal on mutant-mutant interactions.

### The subunits of adaptation

Class-specific inclusive fitness *w*_IF*a*_(*x*, *y*) is the reproductive value of a class-*a* individual augmented by the average effect of that individual switching to expressing a copy of the mutant allele on the reproductive-value weighted number of mutant gene copies produced by all recipients of its action(s). Thus all behavioral effects are grouped by actor in eq. (4) and we emphasize that this holds also within classes (see Supplement B for proofs and for details on the connection to the neighbor-modulated formulation of inclusive fitness). Crucially, inclusive fitness is defined at the allele level and this is consistent with the original formulation of this concept (Hamilton, 1964, pp. 3-8). But our own formulation (eq. 4) extends it to class-structured populations, and further shows that it holds regardless of the complexity of the evolving trait and the strength of selection on the mutant allele.

The fundamental difference between average direct fitness *w*_DF*a*_(*x*, *y*) (eq. 3) and inclusive fitness *w*_IF*a*_(*x*, *y*) (eq. 4) is that the direct fitness is non-null only for individuals who reproduce, while the inclusive fitness effect is non-null only for individuals whose trait affects their own individual fitness and/or that of other individuals in the population (see the concrete social insect example in Box 2). As a result, in an uninvadable population state the inclusive fitness of each class appears to be maximized with respect to the class-specific *x*_*a*_ trait, while the average direct fitness *w*_DF*a*_(*x*, *y*) does not.

Inclusive fitness also makes explicit that a fitness comparison is made between expressing or not the mutant allele, since this involves comparing successful number of gene copies gained and lost through behavioral interactions. Hence, inclusive fitness allows one to fasten attention on the pathways determining fitness costs and benefits (Grafen, 1988). This is particularly salient in the case of classes, where a fitness measure can be attached not only to reproductive individuals but also to, say sterile workers, which can thus be seen as contributing to the fitness of their gene lineage. The inclusive fitness formulation thus shifts attention from those individuals that are passive carriers of alleles to those individuals whose trait actively affects the transmission of alleles. In other words, the unit of adaptation is the gene (Dawkins, 1978, 1982; Haig, 2012), and its subunits are the replicate gene copies expressed differently in particular classes of individuals, whose traits can be regarded as the outcomes of maximizing a class-specific inclusive fitness.

## The rational-actor perspective of adaptation

### Fitness as-if

Class-specific inclusive fitness *w*_IF*a*_(*x*, *y*) (eq. 4) seems satisfying from a population genetics point of view to understand selection on traits affecting relatives. Many behavioral ecologists, however, observe individual behavior instead of genes. What is then missing to understand how selection targets traits in the class-specific representation of inclusive fitness *w*_IF*a*_(*x*, *y*) is a prediction of adaptive traits among the alternatives that an individual can potentially express, *given* the actions of social partners. The simplest and most widely used concept for the prediction of individual behavior is that of a Nash equilibrium trait profile, compared to which no individual can get a higher “payoff” by a unilateral deviation of behavior (see e.g., Luce and Raiffa, 1957, Fudenberg and Tirole, 1991 or Mas-Colell et al., 1995). Here, an individual is envisioned as an autonomous decision-maker, “freely choosing” its action independently of each other individual and with a striving to maximize payoff. In a Nash equilibrium, the individual then makes the best decision for itself in terms of payoff, based on all others individuals making the best decisions for themselves. Our aim now is to find such a payoff for a focal individual of class *a*, denoted 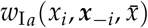. Here we use the same notation as before for individual traits, but *x*_*i*_ is to be understood as any trait value that a given individual *i* could express instead of the (genetically determined) trait that it actually expresses. We refer to 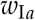 as the *fitness as-if* function of an individual of class *a*, because the individual acts *as if* it maximizes this function in an uninvadable state by choosing the appropriate trait value 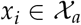 (see eq. C.1 and Supplement C for more details on the construction of fitness as-if).

The previously considered invasion fitness, class-specific average direct fitness, and class-specific inclusive fitness do not fulfill the role of a fitness as-if since they depend on mutant and resident traits. The fundamental difference between fitness as-if *w*_I*a*_ maximization and, say inclusive fitness *w*_IF*a*_ maximization, is that each individual can vary its own trait in fitness as-if, independently of each other individual. By contrast, inclusive fitness is maximized with respect to a mutant trait and this implies that it takes into account correlated changes in the traits expressed by co-bearers. In this respect, the previous perspectives were all gene-centered, while the rational-actor perspective is distinctly individual-centered. In reaching a definition of a fitness as-if, we will still need to take correlated trait changes into account, by adjusting the function definition rather than by allowing its argument ***x***_−*i*_ to vary with *x*_*i*_.

### A general rational actor-centered maximand?

Supplement C shows that it is possible, in theory, to construct an inclusive fitness as-if maximand that individuals appear to maximize in an uninvadable population state (see eq. C.28). This maximand, however, requires that the contextual distributions are written in terms of the actor’s trait and the average in the population. It is thus as if the actor *controlled* the genetic and demographic contexts it experiences. In reality, however, the contextual distributions cannot be under the actor’s control. These distributions do not depend on the actor’s behavior, but on the reproduction and survival of ancestors that determine the present genetic and demographic contexts. This precludes, in our understanding, a general rational actor-centered representation of adaptation.

If the contextual distributions, ***ϕ***(*x*, *y*) and ***q***^D^(*x*, *y*), were to be independent of the mutant trait, then a fitness as-if could be constructed with these distributions exogenous to an individual’s own behavior. There are least two ways to achieve exactly this and both hinge on *weak-selection* approximations implying that, to first-order, the distribution of genetic and class contexts will no longer be dependent on the mutant allele. Such first-order approximations are reached either by assuming that the phenotypic effects of the mutant is small (“small-mutation” weak selection, in which case the individual fitness functions *w*_*ua*_ are assumed differentiable with respect to trait values), or that parameters determining both mutant and resident phenotypic effects on fitness are small (“small-parameter” weak selection). In both cases, the contextual distributions depend at most on the resident trait (***ϕ***(*x*, *y*) *∼ **ϕ***(*y*) and ***q***^D^(*x*, *y*) *∼ **q***^D^(*y*), see Supplement C.5.1 for more details). The key implication is that under weak-selection a mutant allele will not affect the genealogical and/or demographic class structure to which it is exposed and this structure can thus be held constant. This was a central assumption endorsed by Hamilton (1964, p. 34), and has been used to obtain a representation of individual maximizing behavior in the absence of social interactions in age-structured populations (Grafen, 2015; Lessard and Soares, 2016) and in the presence of social interactions in group-structured populations (Lehmann et al., 2015). These two cases can be generalized by the results provided in Supplement C, and we now present a fitness as-if that takes the form of inclusive fitness.

### Maximization of inclusive fitness as-if under weak selection

Assuming weak selection, we show in Supplement C that the inclusive fitness as-if function 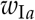 of a class *a* individual defined by

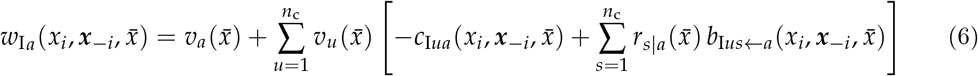

is maximized in an uninvadable population state. In this inclusive fitness as-if, 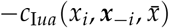 is an average effect on the number of class-*u* offspring produced by a single class-*a* individual and 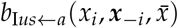 is an average effect of a single class-*a* individual on the number of class-*u* offspring produced by all class-*s* neighbors. These average effects, costs to self and benefits to others, are now weak-selection approximations of the exact costs and benefits obtained by performing a general regression of the individual fitness of *i* when in class *a* on the frequency in itself and its neighbors of a hypothetical allele determining trait expression, whereby the effects of switching trait expression can be assessed. This allele is taken to have the same distribution as the mutant allele in the population-genetic model and ensures that the inclusive fitness as-if (eq. 6) of a class *a* individual aligns, at uninvadability, with the class-specific inclusive fitness of the population genetic model (eq. 4). The key difference between the cost 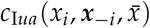 in the as-if and the cost *−c*_*ua*_(*x*, *y*) in the population-genetic model (recall eq. 4) is then that all individuals within groups have distinct traits in the rational-actor perspective (and likewise for the benefits 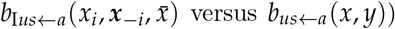 versus *b*_*us←a*_(*x*, *y*)). As such, the probability 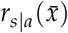 that, conditional on being in class *a*, a random actor and a random class-*s* recipient in its group share the same allele is constant with respect to the actor’s trait. Hence, it is equivalent to the standard relatedness in a monomorphic population with trait 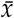 (sometimes called pedigree relatedness) and is independent of actor genotype.

Eq. (6) provides an inclusive-fitness representation of fitness as-if that individuals from each class appear to maximize in an uninvadable population state (see Supplement C.5.2 for a proof of this result). It may be felt to be a stretch to define the inclusive fitness as-if, which is not a population genetic quantity. Such an elaborate construction may nevertheless be needed to assign a coherent meaning to the untold number of statements found in the literature that individuals maximize their own inclusive fitness (rather than the inclusive fitness, eq. (4), of an allele in the population genetic model). Of particular note here is that our construct can be assigned to individuals without knowing their genotype. On the other hand, its representation (eq. 6) still makes explicit that individuals in each class adjusting their behavior will strive to do so by maximizing their genetic contribution to the next generation and by treating others according to the degree of genetic relatedness between actor and recipient.

### Scope of our results

We have identified a rational actor-centered maximand, which individuals appear to maximize in uninvadable population states under weak selection (or if, for other reasons, the ***ϕ*** and ***q***^D^ contextual distributions are independent of selection). This result, as well as the actor-centered maximand (eq. 4), were obtained assuming simplifying demographic assumptions; in particular, constant group size and abiotic environment, no isolation-by-distance, and discrete time. We now confront each of these the assumptions in turn. First, the number of individuals in each class and thus group size as well as abiotic environments are all likely to fluctuate. To cover these cases, it suffices to follow the recommendation of McPeek (2017), which describes common practices in theoretical evolutionary biology (e.g., Brown and Vincent, 1987; Taper and Case, 1992; Geritz et al., 1998; Dercole and Rinaldi, 2008; Lion, 2018), to write individual fitness *w*_*ua*_ not only as a function of an individual’s trait and that of its interaction partners (group and average population members), but also as a function of relevant endogenous variables (e.g., population size, abiotic environment, cultural knowledge); namely, those variables whose distributions or values are influenced by individual traits and thus result in environmental feedbacks on fitness. For weak selection, these distributions or values can then be approximated as a function of the resident traits (see Ronce et al., 2000; Rousset and Ronce, 2004 for concrete examples of fitness functions and distributions covering both demographic and environmental fluctuations, in particular examples where the number of individual in each class can fluctuate between groups, as well as examples where the total number of individuals within groups can fluctuate, respectively). Then, an inclusive fitness as-if under individual control can be defined; the implication being that there are now more contexts to consider relative to the case with no fluctuations (e.g., different group sizes and environments, different number of individuals across classes in different groups), and individuals face a maximization problem under the constraint that the endogenous distributions or values of contexts are evaluated at the uninvadable trait state. Likewise, taking isolation-by-distance into account calls for an extension of the number of contexts and relatednesses to be considered. But given the contexts and the relatednesses, their distributions or values can again be approximated as function of the resident strategies under weak selection (see Rousset and Billiard, 2000 for such constructions for isolation-by-distance) and again an inclusive fitness as-if under individual control can be defined. We also assumed discrete time but comparison of our results (in particular to those of the continuous time model of Grafen (2015, eq. 38) suggests that here again, only a redefinition of contexts is needed to cover fitness as-if under continuous time.

Our results also relied on specific assumptions about trait expression. While traits themselves can be arbitrary complex (e.g., they can be of arbitrary dimension, combining both discrete and continuous trait values, be reaction norms), we assumed that there is no kin recognition or discrimination based on using genetic cues (most models in the literature do not allow for this case either). Yet, other forms of kin discrimination, such as different behaviors expressed toward sisters, cousins, etc., are readily accounted by our results once sisters, cousins, etc., are recognized as different classes of actors. Our model also does not cover complicated conditional class-specific traits expressions, such as when an individual helps its mother as long as she is alive and upon her death starts to help its siblings. Including such biologically relevant cases and more generally conditional trait expression based on individual recognition again calls for an extension of the number of contexts to be considered in the definition of fitness as-if. Finally, we assumed a resident monomorphic population, but this population could be polymorphic with any finite number of genotypes coexisting in equilibrium. Here, previous results (Eshel and Feldman, 1984; Liberman, 1988; Eshel et al., 1998) suggest, again, that an extension of the number of contexts will allow to define a corresponding fitness as-if. In conclusion, all the above scenarios involve considering more complicated ***ϕ*** and ***q***^D^ contextual distributions, so that explicit extensions of our results under these scenarios will need care for the definitions of contexts, fitnesses and relatedness. Such demographic, genetic, and behavioral extensions could be very welcome to generalize both class-specific inclusive fitness (eq. (4)) and class-specific inclusive fitness as-if (eq. (4)), but are unlikely to alter our conclusions, since they all rely only on broadening the number of contexts (be it demographic or genetic).

## Discussion

### Summary and take-home messages

The evolutionary literature provides contrasting messages about the relationship between adaptation and individual behavior as the outcome of fitness maximization. We here combined core elements of evolutionary invasion analysis, inclusive fitness theory, and game theory in order to get a hold on the conditions under which individuals can be envisioned as maximizing their own inclusive fitness in an uninvadable population state (i.e., a population where all deviating mutants go extinct). In particular, we considered three individual-centered perspectives on adaptation (Fig. 1), and defined two class-specific inclusive fitness maximands. The first inclusive fitness maximand is assigned to bearers of a mutant allele, but here only identically to all bearers in a given class, rather than identically to all bearers of an allele as in Hamilton (1964). Our second maximand is assigned to each individual in the population separately, and is a generalisation of our class-specific inclusive fitness, that a behavioral ecologist can directly assign to any observed individual without knowing genotypes. Instead of depending on genotype, it has to depend on the traits expressed by all social partners. This “behavioral” inclusive fitness nevertheless reduces to the class-specific inclusive fitness of a mutant allele when the resident population is an uninvadable population state and under a weak-selection approximation. We thereby defined a rational actor-centered inclusive fitness that is maximized at an evolutionary equilibrium. Our formal analysis leads to the following main take-home messages.

**Figure 1:**
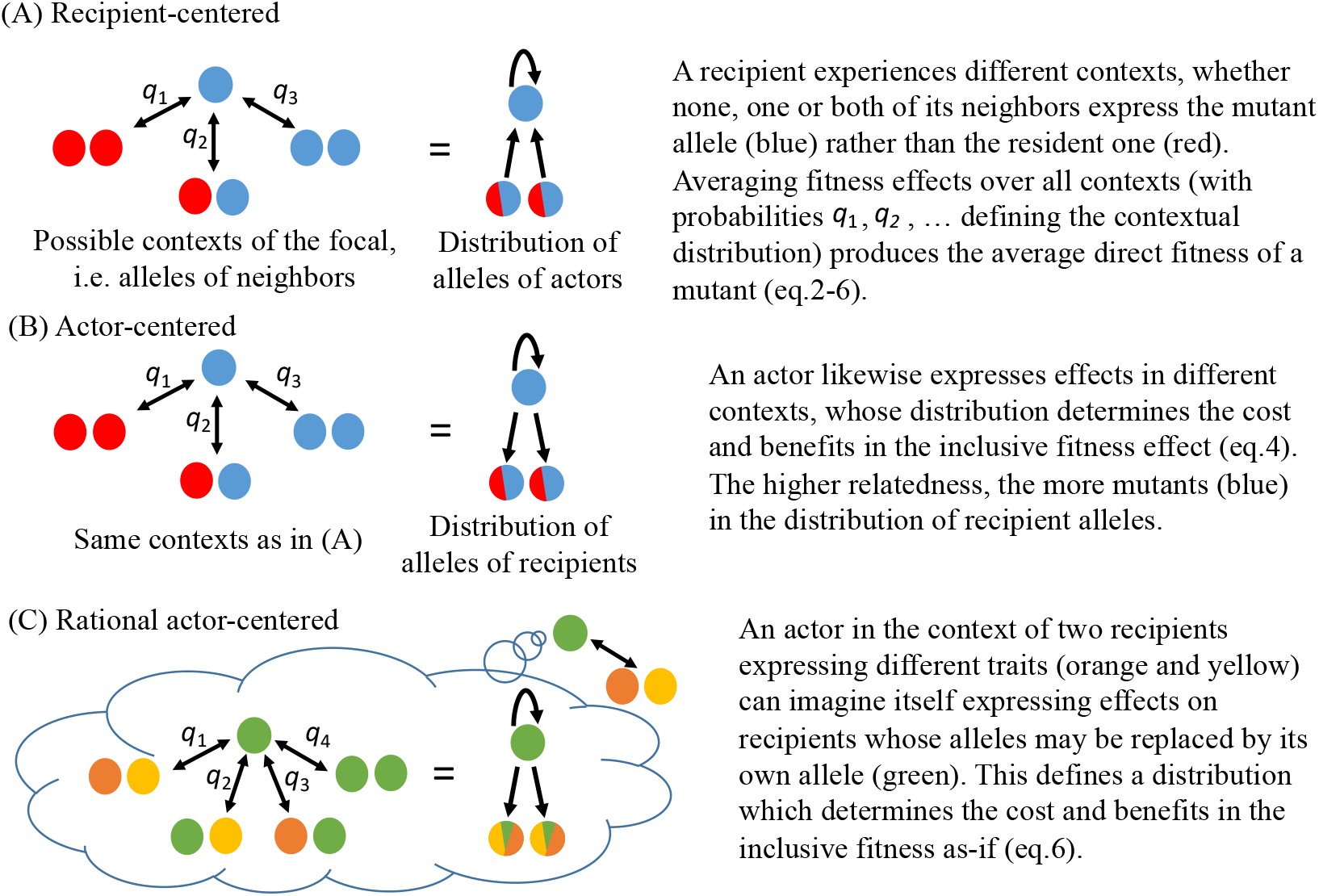
Three perspectives on adaptation. The three perspectives are here illustrated in groups of three haploid individuals, and ignoring any class structure. In the “recipient-centered” perspective (panel A), interactions between group members are described by arrows representing effects of group neighbors on the fitness of a focal recipient carrying a mutant allele. In the “actor-centered” perspective (panel B) and “rational-actor centered” perspective (Panel C), interactions between group members are described by arrows representing effects of a focal actor on the fitness of average group recipients.

**Message (1): actor-centered inclusive fitness maximization obtains generally.** Uninvadable traits can be characterized in terms of mutant alleles attempting to maximize (in the best-response sense) their own transmission across generations. We showed that the traits expressed by individuals in each class maximize class-specific inclusive fitness in an uninvadable population state (eq. 4). This provides a genuine actor-centered perspective of adaptation.
**Message (2): inclusive fitness is a gene-centered fitness measure.** Selection on a mutant allele depends on both the individual fitness of its carriers and the distributions of class and genetic contexts in which these carriers reside. Since these distributions are properties of a lineage of individuals over multiple generations, the class-specific inclusive fitness is not the fitness of a single individual, but that of an average class-specific carrier of a mutant allele sampled from the distributions of genetic contexts it experiences.
**Message (3): rational actor-centered inclusive fitness maximization under weak selection.** For weak selection, the distributions of class and genetic contexts, and thus relatedness, can be taken to be unaffected by selection (Hamilton’s 1964 original modeling assumption). In this case, we showed that each individual in each class appears to maximize its own inclusive fitness (eq. 6) in an uninvadable population state. This provides a genuine individual-centered perspective of adaptation that can be assigned to an individual without knowing its genotype.

Message (1) follows from the fact that alleles are the information carriers of the hereditary components of organismic features and behavior. As emphasized by Dawkins (1979, p. 9), alleles do not act in isolation but in concert with all other alleles in the genome and in interaction with the environment to produce the organism. But uninvadability can be deduced from unilateral deviation of allelic effects alone. This logic can be transposed down at the class level so that adaptation in the long-term evolutionary perspective can be envisioned as the maximization of the class-specific inclusive fitness of a mutant allele that holds for any trait complexity and selection strength. This inclusive fitness consists of the class-specific reproductive value of an allele augmented by the inclusive fitness effect, which is a decomposition of the force of selection, in terms of direct and indirect effects on transmission of replica copies of this allele. The inclusive fitness effect in class-structured populations is commonly represented, to the first-order, as an average effect on the reproductive value weighted-number of offspring produced by a recipient of a given class (Taylor, 1990; Frank, 1998; Rousset, 2004). Our class-specific inclusive fitness, which collects effects by actors of a given class, better embodies the notion of class-specific inclusive fitness and holds generally. Further, since any (indirect) effect that an actor from a given class has on the survival and reproduction of a relative in another class is not an effect on the actor’s own fitness, the maximization of class-specific average direct fitness is generally not compatible with uninvadability, in contrast to class-specific inclusive fitness.

Message (2) follows from the fact that the selection pressure on a social trait depends on what carriers and other individuals are doing. One cannot say what is the best to do for one individual, without specifying the actions of other individuals in present *and* past generations. This applies to inclusive fitness as well, and shows that the fitness effects under the control of an allele (the set of all copies of an allele) may include changes of class (demographic) and genetic contexts. As such, inclusive fitness is a gene-centered fitness measure (consistent with Hamilton’s original definition), whose components, like relatedness, cannot be under the control of a single individual. This precludes a general interpretation of uninvadability as individuals maximizing *their* inclusive fitness. The usual individual-centered characterization of Nash equilibrium in the social sciences (e.g., Luce and Raiffa, 1957; Fudenberg and Tirole, 1991; Binmore, 2007) bears similar limitations as a characterization of human (evolved) behavior.

Message (3) follows from the fact that when selection is weak, the dependence of the inclusive fitness of an allele on the frequency of this allele across generations can be simplified and the outcome of evolution can be regarded as individuals maximizing their own inclusive fitness for a given distribution of genetic-demographic contexts. This warrants the view of individuals from different classes as autonomous decision-makers, each maximizing its own inclusive fitness.

We finally delineate two implications highlighted by our analysis.

### Is it empirically expected that *−c* + *rb >* 0?

First, in any population that is subject to density-dependent regulation and that is in an uninvadable state, the average individual fitness, and the average inclusive fitness, are equal to one. In practice, populations will not be exactly at uninvadable states, but, when they depart from such states, they are not expected to depart in any consistent way in terms of expressing social behavior (e.g., the trait level expressed by individuals could be as well below or above the uninvadable one). Hence, in contrast to the hypothesis considered by Bourke (2014), we do not necessarily expect a tendency for the inclusive fitness effect (“*rb − c*”, the difference between the baseline fitness of “1” and inclusive fitness) of a “more social” act to be positive even when kin selection operates through positive indirect effects (*rb >* 0). This argument applies to the class-specific inclusive fitness as well (the difference between *w*_IF*a*_(*x*, *y*) and the reproductive value in eq. 4, which will be zero at uninvadability).

Bourke (2014) reviewed a number of studies that have attempted to test kin selection by quantifying the inclusive fitness effect, and he documented only a weak tendency for a positive bias. This meta-analysis was framed as a test of Hamilton’s rule, and it could then be seen as providing little support for kin selection theory. But the inclusive fitness effect, *rb − c*, and more generally the class-specific inclusive fitness effect, should be negative for any mutant trait in an uninvadable population state. This is so, since by definition, the inclusive fitness effect is the effect of a mutation away from an uninvadable population state on an invasion fitness maximized at this state. Hence, the results of Bourke’s (2014) meta-analysis could actually be seen as evidence that populations are generally close to some evolutionary equilibrium, where the inclusive fitness effect tends to vanish.

By contrast to the inclusive fitness effect (*rb − c* in the absence of classes), the indirect fitness effect, *rb* will be non-zero when kin selection operates at an evolutionary equilibrium. In testing kin selection theory, a difference should thus be made between attempting to measure the inclusive fitness of an allele, which, at an equilibrium, is not informative about the importance of kin selection, and the indirect fitness effect, which for all conditions quantifies how the force of selection on a mutant depends on relatedness. Nevertheless, the inclusive fitness of a particular class of individuals (eq. 4) is itself informative about the importance of kin selection, since it assigns fitness contributions even to individuals that do not reproduce.

### So, when do individuals maximize their inclusive fitness?

The second and more significant implication of our results is to support the common conception in behavioral ecology and evolutionary psychology, of adaptation as the result of interacting individuals maximizing their own inclusive fitness (e.g., Alexander, 1979, 1990; Alcock, 2005; Buss, 2005; Grafen, 2007, 2008; Davies et al., 2012; West and Gardner, 2013; Crespi, 2014). Insofar as evolutionists think about adaptation in this way, they should keep in mind the underlying weak-selection assumption (points 2 and 3), the assumption that the population needs to be close to an uninvadable state, and, the definition of inclusive fitness for which it holds (eq. 6). Namely, it is a function of the traits of an individual and of its social partners all assumed distinct, yet which coincides with the allelic inclusive fitness of an allele at an evolutionary equilibrium. Previous work has been able to justify that individuals appear to be to maximize their inclusive fitness only for behaviors that do not involve any phenotypic interactions (Grafen, 2006a)^1^, whereby ruling out social interactions in any broad sense.

While Fisher (1930) and Hamilton (1996, p.27–28) have emphasized the importance of weak selection for the evolutionary process, weak-selection approximations are still sometimes vilified in evolutionary biology (as reviewed by Birch, 2017). The value of approximations, however, can only be assessed by their impact on a field. Humans were landed on the Moon using Newtonian mechanics (Wakker, 2015)–a first-order approximation to the real (relativistic) mechanics of the solar system (Okun, 2012). Thus, technological and scientific achievements regarded as paradigmatic are as dependent on approximations as is the individual-centered version of inclusive fitness. A number of unique predictions about social behavior have been made by focusing on individual inclusive fitness-maximizing behavior, from conflicts over sex-ratios and resources within families to inbreeding tolerance and genomic imprinting (e.g., Trivers and Hare, 1976; Haig, 1997a; Alcock, 2005; Macke et al., 2011; Davies et al., 2012; Szulkin et al., 2013). Our analysis justifies formally this long-heralded view.

## Acknowledgments

We thank R. Bshary for having initially stimulated the writing of this paper and T. Priklopil for useful discussions. We also thank M. Chapuisat and N. Raihani for useful comments on previous version of the manuscript. We gratefully acknowledge extensive and extremely useful comments on the paper by D. Bolnick, A. Grafen, S. Lion and an anonymous reviewer. These comments fundamentally improved the manuscript, the results as well as the presentation.

---

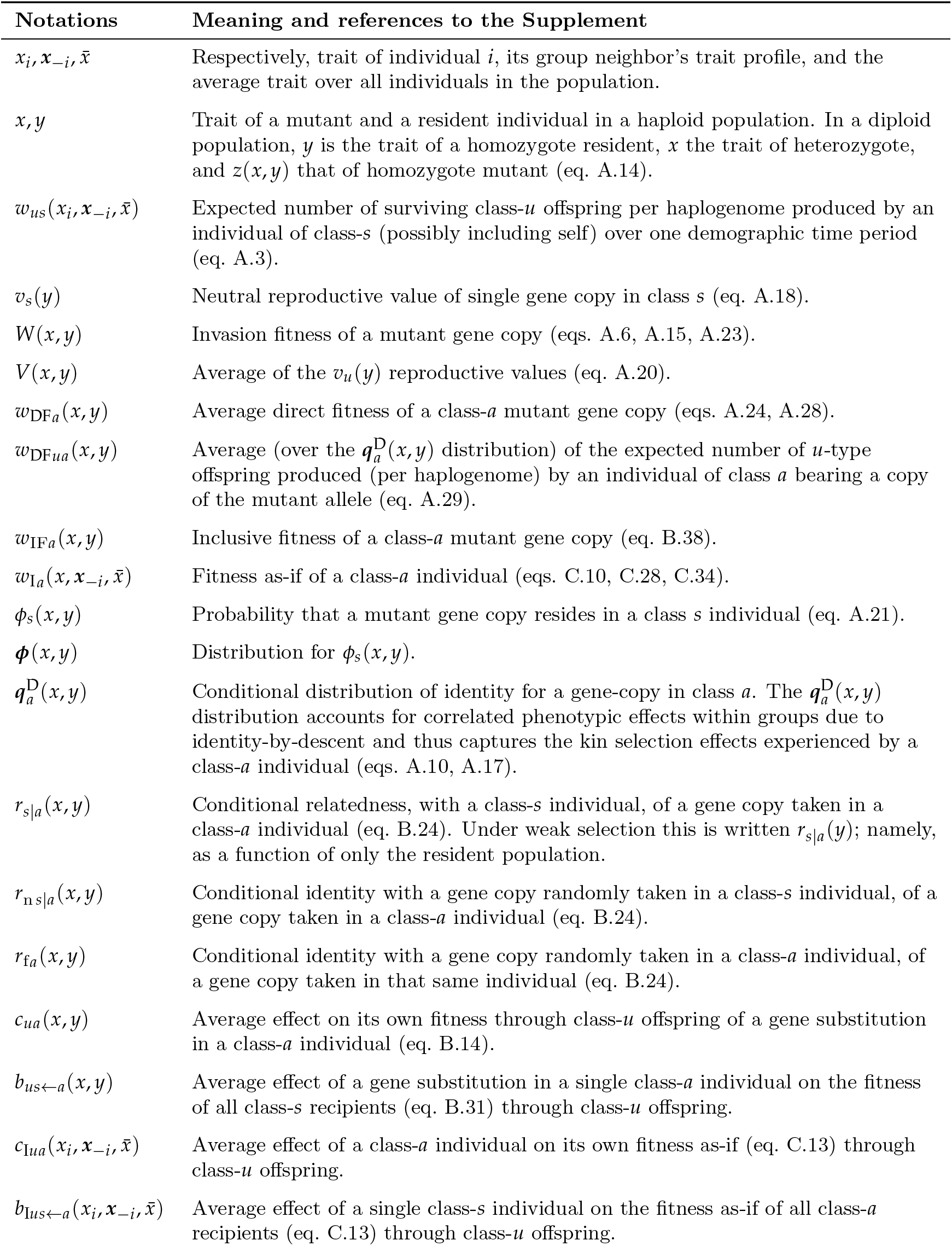

## Box 1: Social insect example

Let us thus consider a seasonal population of diploid social insects (e.g., termites rather than ants) who allocate resources to the production of three classes of individuals, reproductive males, reproductive females (queens), and workers. The life cycle is as follows. (1) At the beginning of the season each group is occupied by exactly a single mated queen that initiates a colony by producing workers that help produce sexuals. (2) At the end of the season, all reproductive individuals disperse at the same time and individuals of the parental generation die. (3) Random mating occurs, all queens mate exactly with one male and then compete for vacated breeding slots to form the next generation. Under these assumptions, successful gene copies must pass through the single mated female in each group; and the expected number of class-*u* offspring produced by a class-*a* mutant individual per haplogenome can be written *w*_DF*ua*_(*x*, *y*), where trait *x* = (*x*_f_, *x*_m_, *x*_o_) collects, respectively, the traits of females, males, and workers. The trait for the resident is *y* = (*y*_f_, *y*_m_, *y*_o_), whereby the invasion fitness of the mutant can be written as in eq. (2) with the direct fitnesses given by

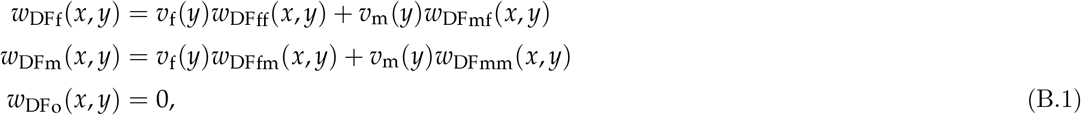

and the distribution of class states in which a carrier of the mutant allele can reside being ***ϕ***(*x*, *y*) = (*ϕ*_f_(*x*, *y*), *ϕ*_m_(*x*, *y*), *ϕ*_o_(*x*, *y*)). A worker does not reproduce and henceforth has a zero direct fitness, but its class frequency is non-zero (*ϕ*_o_(*x*, *y*) ≠ 0; the explicit expression for *ϕ*_f_(*x*, *y*), *ϕ*_m_(*x*, *y*), and *ϕ*_o_(*x*, *y*), are given in the Supplement, eq. A.32) and it helps its parents to reproduce. Formally, the worker affects the reproduction of its male and female parents through the dependence of individual fitness on trait vector 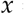 = (*x*_f_, *x*_m_, *x*_o_).

## Box 2: Individual and inclusive fitness for social insect example

As an example of the individual fitnesses in eq. (B.1), making the (evolutionary) role of workers more explicit, let us assume that each female produces exactly one worker, which increases colony productivity according to its trait *x*_o_. We also consider that the female trait *x*_f_ determines the sex-ratio, and nothing else. Finally, we consider that the male trait does not affect any fitness component. Since the worker is heterozygote with probability 1/2 and homozygote for the resident with probability 1/2, the expected number of (reproductive) daughters of a mutant female can be written as

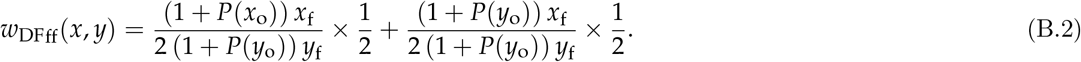

Here, *x*_f_ is the proportion of offspring that become female. The worker affects the relative fecundity of a female, which is assumed to be given by 1 + *P*(*⋅*), where *P*(*⋅*) is some function of worker trait. In other words, the worker trait increases offspring production of the queen relative to some baseline. The first term in eq. (B.2) is for the case where the worker is heterozygote and the second when it is homozygote resident. The denominators in eq. (B.2) reflects female production by other colonies that are monomorphic for the resident allele and the 2 reflects the fact that we measure fitness per haplogenome. Likewise, the number of sons produced by a female is

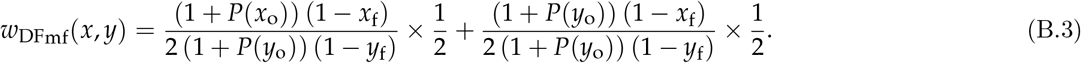

While male trait does not impact fitness, the mutant allele may still occur in a male and the mutant male fitness components will depend on the worker trait, which affects offspring production by the male’s mate(s), whereby

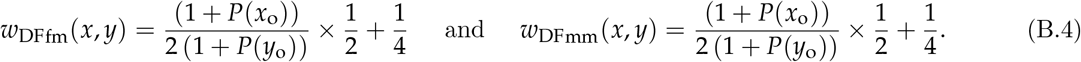

For this model, the inclusive fitness effects of a female, male, and worker carrying the mutant in a population at the equilibrium sex-ratio of 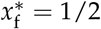 are, respectively,

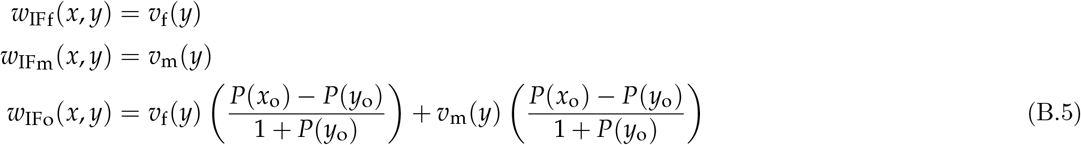

(see Supplement B.3 for a proof).

## Supplement A: Invasion fitness as average direct fitness

In this Supplement, we derive the expressions for the average direct fitness given in the main text, eqs. (2)–(3). As explained therein, we limit our discussion to a population that is divided into an infinite number of groups that are all of constant size *n* and connected by random dispersal with reproduction occurring in discrete time periods (i.e., Wright’s 1931 canonical island model of dispersal). In each group, there is a finite number of classes and we use the following notations (see also section “Demographic assumptions” of the main text): *n*_*a*_ denotes the number of individuals in class *a*, *C* denotes the set of classes (e.g., workers and queens, males and females; 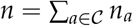 and 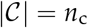), and 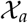 denotes the feasible trait set of an individual of class 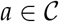 with the total trait set being 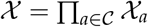 (products of sets are taken as Cartesian products throughout).

In such a population, a resident trait *x*^∗^ is uninvadable if it is a best response to itself, meaning that if we can vary invasion fitness *W*(*x*, *x*^∗^) by varying the mutant trait *x* in the set 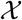 of feasible traits,^2^ an uninvadable trait must be a trait maximizing invasion fitness for the resident at *x*^∗^ (formally invasion fitness is the function 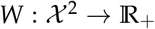 with *W*(*y*, *y*) = 1 for all 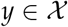). It then follows that an uninvadable trait *x*^∗^ satisfies

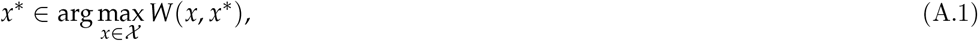

which means that *x*^∗^ belongs to the set of traits resulting in the highest invasion fitness among all alternatives given in the set 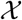 of feasible traits, for the resident population at *x*^∗^ (in eq. A.1, *x*^∗^ belongs to a set because an uninvadable trait is not necessarily unique).

Since (even assuming the island model) notations and concepts, to express the function *W* in terms of individual-centered components in the presence of class-structure and diploidy, become rapidly complicated, we will progressively introduce different cases (haploid, diploid, etc.), concepts, and notations. We start by defining formally the central building block of our analysis, which is individual fitness.

### A.1 Building blocks

#### A.1.1 Individual fitness

In the absence of class-structure, we define the *individual fitness function* as

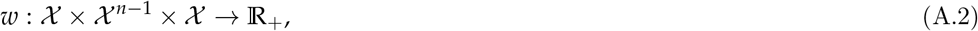

such that 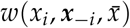 is the expected number of successful offspring produced (*per haplogenome*) by a focal individual 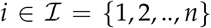 with trait 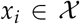, where neighbors in group 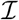 have trait profile 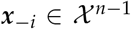 in a population where the average trait over all individuals is 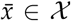.

The expectation is over all within-generation stochastic effects on settled offspring number in the descendant generation and conditional on realized trait profile 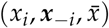 in the parental generation.

In the presence of class-structure, and following the assumptions and explanations of the main text, the individual fitness functions is defined as

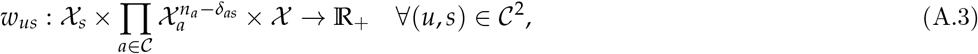

such that 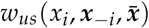 is the expected number of successful class-*u* offspring produced over a demographic time step by individual 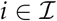 when in class-*s* (*per haplogenome*) with trait 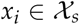 in a group where neighbors have trait profile 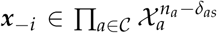 (which has dimension *n −* 1 owing to the fact that *δ*_*as*_ is the Kronecker delta) in a population where the vector of average traits is 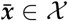.

#### A.1.2 Distinct and indistinct individuals

The formulation of the fitness functions (eqs. A.2–A.3) allows for a characterization of the population where each individual in a group can be distinguished from each other. This means that the trait profile (*x*_*i*_, ***x**_−i_*) in a focal group, i.e., its state, belongs to the set of all ordered trait profiles (i.e., all ordered group states are considered; for instance, in the absence of class-structure, this is 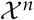). In an evolutionary invasion analysis, however, we consider that only two alleles –mutant and resident– segregate in the population and so there can be a maximum number of only two types of individuals in each class in a haploid population (or three types in diploids: one heterozygote and the two homozygotes). Hence, we have group states with *n* individuals, where each member belongs only to one among a finite number of genotypic types. This allows for an alternative characterization of the population, where one counts the number of individuals bearing identical traits in a group and thus individuals are no longer distinguished (i.e., only unordered groups states are considered).

To illustrate these concepts, consider a haploid population without class structure with individuals either expressing a mutant trait 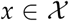 or expressing a resident trait 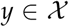. Since each individual in a group is either mutant or resident, there is a total number of 2^*n*^ ordered groups states. Since neighbors are exchangeable, the individual fitness of an individual *i* with given trait *x*_*i*_ is identical for all permutations of neighbors’ traits in ***x***_−*i*_. Thus,

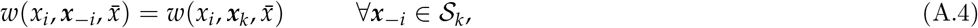

where ***x***_*k*_ = (*x*, *x*, … *y*, *y*, …) is the vector of dimension *n −* 1 with the first *k −* 1 entries equal to *x* and the subsequent *n − k* entries equal to *y*, and 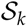 is the set of all distinct permutations of ***x***_*k*_. The number of distinct permutations is the binomial coefficient

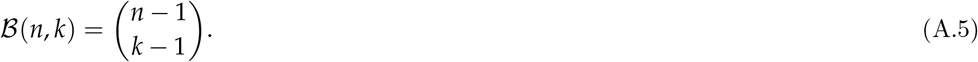

In the class-structured case, permutation-invariance is on the trait profile of neighbors belonging to the same class. Permutation-invariance is not an assumption, it is part of what allows one to determine whether different neighbors belong to the same class of actors. When permutation-invariance does not hold, individuals belong to different classes.

These considerations show that one can characterize a group state in a class-structured population from the perspective of an individual *i* either by distinguishing all individuals (ordered group states) or by not distinguishing individuals in identical states (unordered group states). While in evolutionary analysis individuals are usually not distinguished because this is often mathematically simpler (an exception being the Price equation, Price, 1970; Frank, 1998), distinguishing them is fundamental to the rational actor-centered perspective of adaptation. As such, we develop the invasion fitness by distinguishing individuals when this will be needed for the analysis of the indivdual-centered perspective, but start by not distinguishing individuals to frame the model into the classical approach and to introduce concepts in a progressive way.

### A.2 Average direct fitness without classes

#### A.2.1 Haploids

##### Indistinct individuals

In the absence of within-group class structure (homogeneous individuals), the invasion fitness can be written

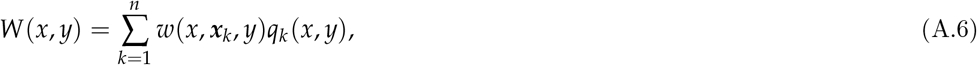

which takes the form of average direct fitness. Indeed, here *w*(*x*, ***x***_*k*_, *y*) is the individual fitness given by eq. (A.4) and *q*_*k*_(*x*, *y*) is the probability that a randomly sampled mutant individual from the mutant lineage descending from the initial mutant resides in a group with *k* mutants 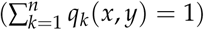.

The *q*_*k*_(*x*, *y*) probability is evaluated under the assumption that the mutant is overall rare in the population, and that the growth of the mutant lineage descending from a single initial copy has reached stationarity. That is,

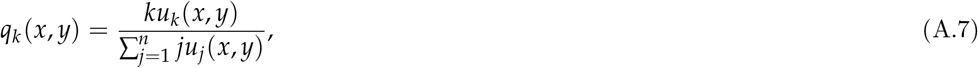

where ***u*** = (*u*_1_, *u*_2_, …, *u*_*n*_) is the right eigenvector associated to the leading eigenvalue *W*(*x*, *y*) of the matrix ***A***(*x*, *y*) describing the growth of the mutant when it is overall rare in the population:

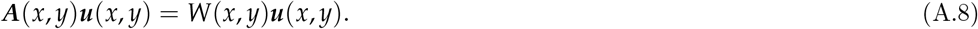

The *jk*th entry of ***A***(*x*, *y*), denoted by *a*_*jk*_, gives the expected number of groups with *j ≥* 1 mutants descending over one time step from a group with *k ≥* 1 mutant, and *u*_*j*_(*x*, *y*) is the stationary probability that there are *j* mutants in a group, conditional on there being at least one mutant. A proof of eq. (A.6) follows by left-multiplying eq. (A.8) with a vector **n** whose entry *j* is equal to the number *j* of mutants, noting that 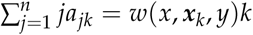 is the expected number of successfull mutant copies produced over one time step by all mutant gene copies in a group in state *k*, and rearranging terms; see Lehmann et al. (2016) for a detailed proof and a more detailed characterization of the multitype branching process underlying mutant dynamics.

##### Distinct individuals

We now make the link to characterizing invasion fitness by considering all ordered groups states. To obtain this representation, we note that from eq. (A.4), we can write

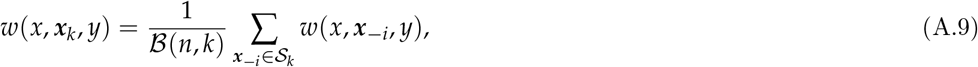

where on the right-hand side we have distinguished all trait profiles in the focal group with *k −* 1 neighbors bearing the mutant allele. Let us now further define

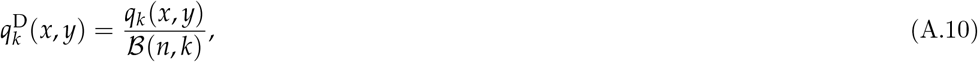

which is the probability that, conditional on an individual carrying the mutant allele, an ordered neighbor trait profile 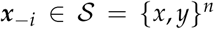 contains exactly *k −* 1 individuals also carrying the mutant (hence 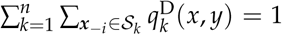 and the superscript D stands for a reminder that the distribution is over profiles of traits for distinct individuals). On substituting eqs. (A.9)–(A.10) into eq. (A.6), we can write the invasion fitness of a mutant allele with trait *x* introduced into a haploid resident population with trait *y* as

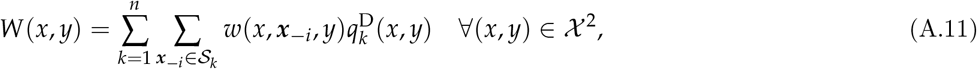

which is the average fitness over all ordered trait profiles in a group.

Writing explicitly the sums appearing in eq. (A.11) and detailing the permutation under the more general diploid and class-structured model will be cumbersome and we now present an alternative and more compact representation of invasion fitness. To that end, let us collect all 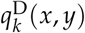 probabilities into the vector ***q***^D^(*x*, *y*), which is the distribution of ordered group states experienced by an individual with trait *x* and that has sample space in 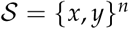. With this, we can write invasion fitness as

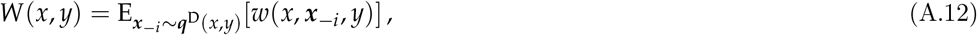

where the notation *∼* specifies that variable ***x***_−*i*_ follows distribution ***q***^D^(*x*, *y*).

#### A.2.2 Diploids

When individuals are diploid, we need to take into account that they can be homozygote for the mutant allele. To do this, it will be convenient to build on our notations for mutant and resident traits introduced for a haploid population. For a diploid population, we let 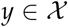 be the trait of an individual that is homozygote for the resident allele and 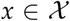 be the trait of an individual that is heterozygote for the mutant allele. Let 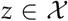 denote the trait of a homozygote mutant and assume that the trait of an heterozygote is obtained as the following convex combination of the trait of the two homozygotes:

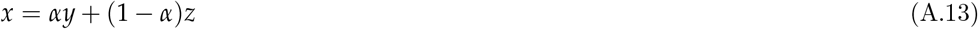

for the scalar *α ∈* (0, 1). Hence, we rule out over-, under-, and strict dominance, but otherwise allow for arbitrary gene action. eq. (A.13) guarantees that for all 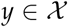 and 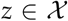, we have 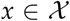 and it allows us to express conveniently the trait of a homozygote mutant as a function 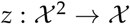 of heterozygote and resident homozygote traits, where

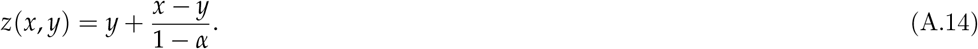

For arbitrary group size *n*, the invasion fitness of a mutant allele with heterozygote trait *x* introduced into a resident diploid population with homozygote trait *y* can be written as

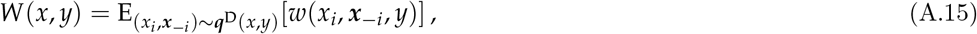

where *x*_*i*_ ∈ {*z*(*x*, *y*), *x*}. Each component *x*_*j*_ of the neighbor trait profile ***x***_−*i*_ = (*x*_1_,, …, *x_i−_*_1_, *x*_*i*+1_, …*x*_*N*_) takes values in the set {*z*(*x*, *y*), *x*, *y*}. The expectation in eq. (A.15) is over the distribution ***q***^D^(*x*, *y*) of ordered group profiles of traits, determined by the distribution of contexts of copies of the mutant allele. The sample space of the distribution of ordered group profiles of strategies is 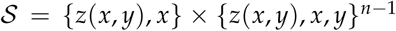. eq. (A.15) shows that invasion fitness can, as in the haploid case and regardless of the mating system, be expressed as an average of the direct fitness components *w*(*⋅*, *⋅*, *⋅*) over a distribution ***q***^D^(*x*, *y*), but which is generally more involved than in the haploid case. Indeed, for the diploid case, the matrix ***A***(*x*, *y*) defining the closed “dynamically sufficient” system determining the fate of the mutant when rare (recall eq. A.8) has entries *a*_**jk**_, which gives the expected number of groups in state **j** descending over one time step from a group in state **k**, and where a state specifies the number of homozygote and heterozygote individuals in a group. More specifically, **j** = (*j*_*y*_, *j*_*x*_, *j*_*z*_), where *j*_*y*_ denotes the number of homozygote residents, *j*_*x*_ the number of heterozygotes, and *j*_*z*_ the number of homozygote mutants in a group. The state space of the process is *𝒢* = {**j**: *j*_*y*_ + *j*_*x*_ + *j*_*z*_ = *n* and *j*_*x*_ + *j*_*z*_ > 0} and the stationary distribution satisfying eq. (A.8) is ***u***(*x*, *y*) = (*u*_**j**_(*x*, *y*))_**j***∈𝒢*_.

A proof of eq. (A.15) then follows by left-multiplying eq. (A.8) with a vector **n** whose entry **j** is equal to the number *j*_*x*_ + 2*j*_*z*_ of mutant gene copies in that state and rearranging terms by noting that ∑_**j***∈𝒢*_ (*j*_*x*_ + 2*j*_*z*_) *a*_**jk**_ = *w*(*x*, ***x***_k_, *y*)*k*_*x*_ + *w*(*z*(*x*, *y*), ***x***_k_, *y*)2*k*_*z*_ is the expected number of successful mutant copies produced over one demographic time step by all mutant gene copies in a group in state **k**, and where ***x***_k_ denotes the vector of neighbor group profiles when a group is in state **k**. With this, 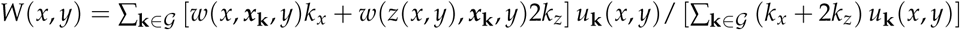 regardless of the assumptions on the mating system. Expanding therein the individual fitnesses in terms of ordered group profiles (like in eq. A.9) one obtains eq. (A.15). The distribution ***q***^D^(*x*, *y*) of ordered group states experienced by a typical copy of the mutant allele will thus be expressed in terms of ***u***(*x*, *y*), of the numbers *k*_*x*_ and 2*k*_*z*_, and combinatorial terms. We skip the explicit expression of ***q***^D^(*x*, *y*) since it requires to define notations to take into account that a mutant allele copy can be in a homozygote or an heterozygote individual (see eq. A.16 for an example) but the logic to obtain ***q***^D^(*x*, *y*) is as in the haploid case (see eq. A.7 and eq. A.10).

As an example of eq. (A.15), let us consider the case *n* = 2, then

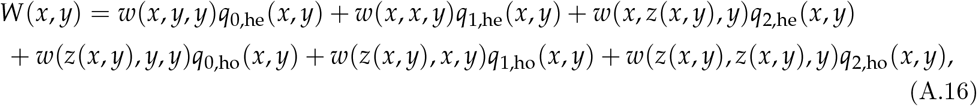

where *q*_*j*,he_(*x*, *y*) is the probability that, given a gene copy is mutant, its carrier is heterozygote and its group neighbor has *j* copies of the mutant (*j* = 0, 1, 2, then indicate the cases, respectively, for the neighbor to be homozygote resident, heterozygote, and homozygote mutant), while *q*_*j*,ho_(*x*, *y*) is the probability that, given a gene copy is mutant, its carrier is homozygote and its group neighbor has *j* copies of the mutant. For this case, the distribution over group contexts is given by

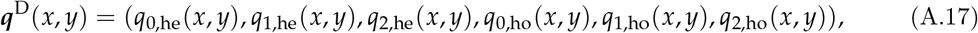

whose elements sum up to one and could be expressed in terms of probabilities of identity in state of alleles in pairs of individuals (Michod, 1982, Fig. 1).

### A.3 Average direct fitness with classes

#### A.3.1 Haploids

In the presence of classes, the trait *x* of the mutant in a haploid population is taken as a vector of actions (or stream of actions), one for each class the individual may belong to, so we write 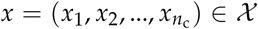, where *x*_*a*_ is the trait of a mutant individual when of class *a*. Likewise, we have 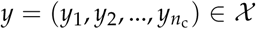. Using eq. (A.3), we let *w*_*us*_(*x*_*s*_, ***x***_k_, *y*) be the expected number of class-*u* offspring produced by a class-*s* mutant when in a group in state **k** = (*k*_1_, …, *k*_*nc*_), which is the vector of the number of individuals carrying the mutant allele in each class, with *k*_*a*_ being the number of mutants in class *a*, whereby ***x***_k_ is a vector that has (*k*_*s*_ − 1) entries with trait *x*_*s*_, *k*_*a*_ entries with trait *x*_*a*_ for each *a* ≠ *s*, while all remaining entries are for the corresponding element of the resident trait vector *y*.

A central quantity in our analysis is the reproductive value *v*_*s*_(*y*) of a single gene copy residing in an individual of class *s* in a monomorphic resident population (neutral reproductive value), which satisfies

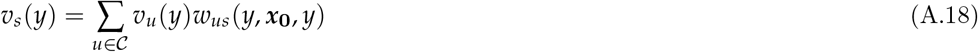

(e.g., Taylor, 1990; Frank, 1998; Rousset, 2004; Grafen, 2006b; Lehmann et al., 2016), where **0** indicates the value of **k** with no mutants at all. With these definitions, the invasion fitness of a mutant allele with trait *x* in a resident population with trait *y* can be written as a sum over, respectively, possible group states, offspring classes, and parent classes:

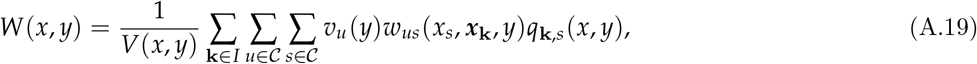

where *q*_**k**,*s*_(*x*, *y*) is the probability that a randomly sampled member of the mutant lineage finds itself in class *s* and in a group in state **k**; 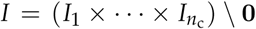 is the set of possible group states with *I*_*u*_ = {0, 1, …, *n*_*u*_} being the set of the number of mutant alleles in class *u*; and

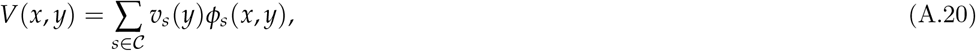

where

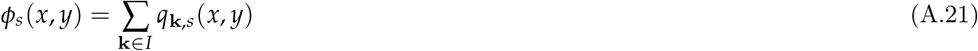

is the probability that a randomly sampled gene copy from the mutant lineage resides in a class-*s* individual. Hence, *V*(*x*, *y*) is the total (neutral) reproductive value of a randomly sampled mutant gene copy from its lineage. Owing to eq. (A.18), *V*(*x*, *y*) can be seen as the average reproductive value of a mutant gene copy that would have its fitness components assigned those of a resident copy (instead of expressing mutant fitness components, the *w*_*us*_(*x*_*s*_, ***x***_k_, *y*)’s, it expresses resident fitness components, the *w*_*us*_(*y*_*s*_, ***x***_0_, *y*)’s).

Eq. (A.19) is in terms of unordered neighbor profiles charaterized by **k**. In this formalism, invasion fitness *W*(*x*, *y*) still satisfies eq. (A.8), if the matrix ***A***(*x*, *y*), describing the growth of the mutant lineage when rare in the population, now has entries *a*_**jk**_ giving the expected number of mutant copies in context **j** that descend from a mutant copy in context **k**, and the *q*_**k**,*s*_(*x*, *y*) distribution is then expressed in terms of the leading right eigenvector ***u***(*x*, *y*) = (*u*_**k**_(*x*, *y*))_**k***∈I*_ of this matrix; namely,

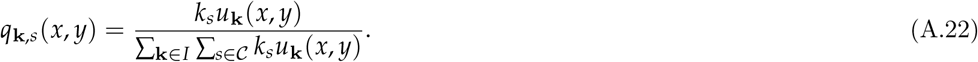

A detailed proof of eq. (A.19) can be found in Lehmann et al., 2016, Appendix F. It follows by left-multiplying eq. (A.8) with a vector **n** whose entry **j** is equal to the reproductive value-weighted number 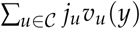 of mutant gene copies in context **j**. This yields 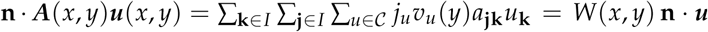, wherein the central expression can be simplified (and the whole expression rearranged) by noting that 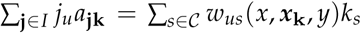 is the expected number of successful mutant copies in class *u* produced over one demographic time step by all mutant gene copies in a group in state **k**.

We now write eq. (A.19) as

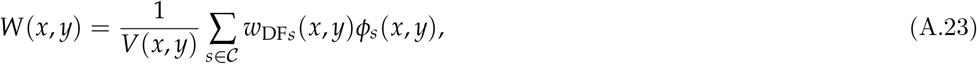

where

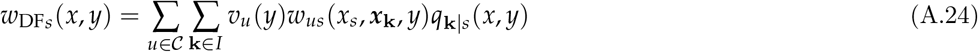

is the expected reproductive value-weighted fitness of a class *s* mutant gene copy and

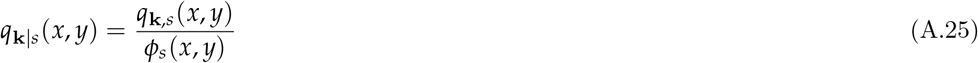

is the probability that, conditional on an individual carrying the mutant allele and of class *s*, the individual resides in a group in state **k**. Eq. (A.24) is the sum of the reproductive values of the descendants of an individual of class *s*, including its potentially surviving self. We thus refer to *w*_DF*s*_(*x*, *y*) as the average direct fitness of a class *s* individual, and, for a panmictic population, this quantity was previously called Williams’ reproductive value (Grafen, 2015, p. 8). Hence, invasion fitness eq. (A.23) is the total average direct fitness of a mutant relative to the reproductive value that individual would have if it expressed the resident trait.

However, any non-null vector of weights could have been chosen in eq. (A.19) and eq. (A.23) to compute the geometric growth rate, which is so because the right-hand side eq. (A.19) is obtained by rearranging the leading eigenvalue-eigenvector equation, where the leading eigenvector can be normalized by any non-null vector (see Lehmann et al., 2016, Appendix B and C for more details). We can in particular choose the unit vector (1, 1, …, 1), whereby invasion fitness becomes the average of the individual fitnesses of a randomly sampled mutant from its lineage. In eq. (A.19), we choose reproductive-value weights for two reasons. First, average direct fitness is then expressed with the same weights as is inclusive fitness (see next section “Inclusive fitness”), given that for inclusive fitness there is no choice but to use the reproductive-value weights. Second, the reproductive-value weights play a pivotal role in the forthcoming weak-selection analysis (section “Individual maximands under weak selection”), where they allow to obtain meaningful expressions for the different average fitnesses, a feature that follows from the well-established fact that the reproductive-value weights are also the unique weights that would allow to apply eqs. (A.23) when the mutant is no longer rare to predict the direction of average allele frequency change by a scalar fitness measure at all allele frequencies under weak selection (e.g., Rousset, 2004; Grafen, 2006b).

Finally, we note that we could normalize the reproductive values such that *V*(*x*, *y*) = 1, however this would induce the *v*_*u*_(*y*)’s to become a function of the mutant, since the *ϕ_s_*(*µ*, *y*) probabilities in eq. (A.20) depend on the mutant. We would like to avoid this here, otherwise differentiation of *W*(*x*, *y*) requires differentiating the reproductive values, and so we need a notation distinguishing the case where reproductive values depend on the mutant from the case of weak selection (investigated below), where this dependence drops out. In order to have a uniform notation throughout the main text, we normalize the *v*_*u*_(*y*)’s such that the average neutral reproductive value of a randomly sampled resident individual is one:

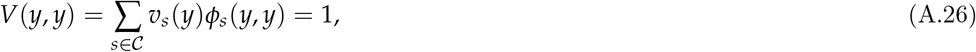

where *v*_*s*_(*y*)*ϕ*_*s*_(*y*, *y*) can be recognized as the reproductive value of class *s* in a monomorphic resident population (e.g., Taylor, 1990; Rousset, 2004) and so eq. (A.26) is the standard normalization of the reproductive values.

#### A.3.2 Diploids and social insects

In order to generalize eq. (A.24) to diploidy, we let 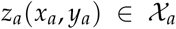 be the trait of an homozygote mutant of class *a* when the profile of heterozygote mutant traits across classes is 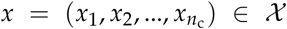 and the trait profile of a homozygote resident individual is 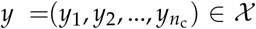(following eqs. A.13–A.14, we assume that for each *a*, *z*_*a*_(*x*_*a*_, *y*_*a*_) is obtained by assuming that heterozygote traits are a convex combination of the homozygotes’ traits). With this, let 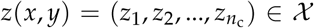 denote the profile of homozygote mutants. Then, the invasion fitness of a mutant allele with heterozygote (multidimensional) trait *x* introduced into a resident diploid population with homozygote trait *y* can be written as

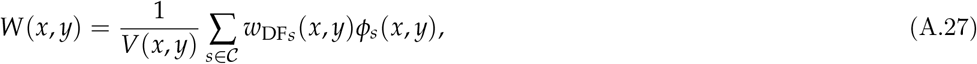

where

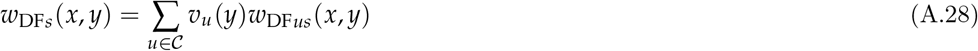

and

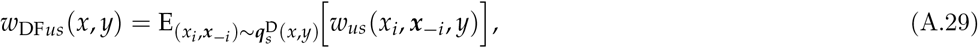

which is the expected reproductive value-weighted fitness of a class *s* mutant gene copy. Here, *x*_*i*_ ∈ {*z*_*s*_(*x*_*s*_, *y*_*s*_), *x*_*s*_} if individual *i* is of class *s* and each component *x*_*j*_ of the neighbor trait profile ***x***_−*i*_ = (*x*_1_, …, *x*_*i−1*_, *x*_*i*+1_, …*x*_*N*_) takes values in {*z*_*a*_(*x*_*a*_, *y*_*a*_), *x*_*a*_, *y*_*a*_} if the corresponding individual *j* is of class *a*. In eq. (A.29), the couple (*x*_*i*_, ***x***_−*i*_) follows the distribution 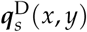 of ordered focal group trait profiles determined by the distribution of contexts of copies of the mutant allele in individuals of class *s*. The distribution of trait profiles has sample space

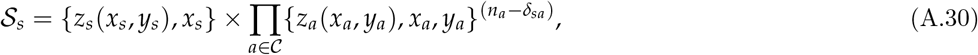

since among the neighbors of an individual of class *s* we have *n*_*a*_ individuals of class *a* ≠ *s* and *n_s_ −* 1 class-*s* individuals. An algebraic proof of eqs. (A.27)–(A.29) follows by combining the line of arguments used to derive the invasion fitness for diploids without classes (eq. A.15) and that for haploids with classes (eq. A.19).

A special case of eq. (A.29) is when there is only a single adult individual per group under complete dispersal and random mating. In that case, a mutant individual can only be heterozygote (as long as the mutant is rare). As a concrete example, we work out the model of a seasonal population of social insects presented in Box 1, where *x* = (*x*_f_, *x*_m_, *x*_o_), which collects, respectively, the traits of females, males, and workers so that the set of classes is 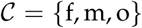. Since we are interested in considering the three classes of individuals demographically, the census stage of fitness is taken right before dispersal (end of stage (1) of the life cycle). When the mutant allele is rare, the dynamics of the number of mutant allele copies in females, males, and workers in the population between successive census stages can be described by the matrix

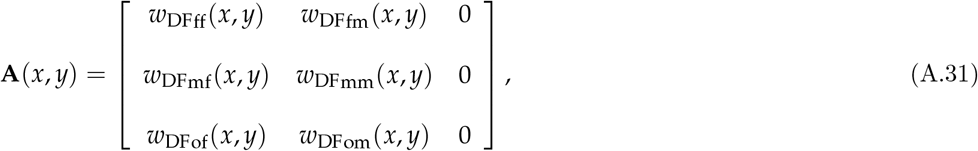

From this matrix, the probabilities that a randomly sampled copy of the mutant allele is in a female, male, or worker, are respectively

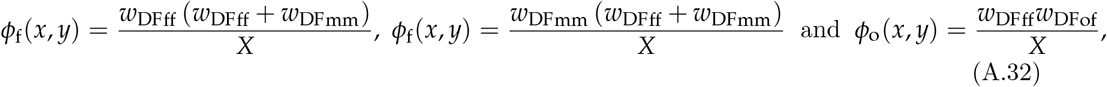

where 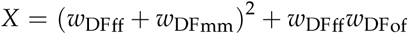. The reproductive values are *v*_o_(*y*) = 0, *v*_f_(*y*) *>* 0 and *v*_m_(*y*) *>* 0, and the invasion fitness is given by

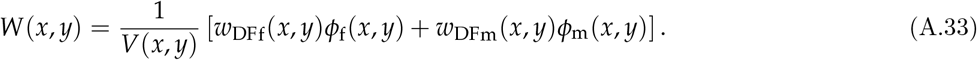

The two direct fitnesses appearing in this equation are given by eq. (B.1) of the main text. Supposing there is only one worker in the colony (e.g., assumptions in the main text), then, in a monomorphic population, we have *ϕ*_f_(*y*, *y*) = *ϕ*_m_(*y*, *y*) = *ϕ*_o_(*y*, *y*) = 1/3 and the reproductive values, normalized so as to satisfy eq. (A.26), are

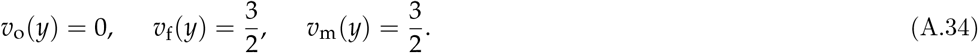

## Supplement B: Inclusive fitness

The main aim of this Supplement is to derive from invasion fitness the expression for the inclusive fitness of a class-*a* individual given in the main text (eq. 4) and then to show that it is maximized for the class-specific trait *x*_*a*_ in an uninvadable state *x*^∗^. As in the previous section, we do so by progressively introducing the different concepts. Before delving into the calculations, it is worth recalling that “inclusive fitness” can be regarded as achieving two related decompositions of the force of selection on a mutant allele. First, it is a partition of selection into direct and indirect fitness effects (the “cost” and “benefit” of Hamilton’s rule, e.g., Hamilton, 1964; Frank, 1998; Rousset, 2004), where the indirect effects are weighted by relatedness coefficient(s). This allows to usefully classify behaviors in different categories (“selfishness”, “altruism”, “spite”, etc.) and is most easily achieved in an evolutionary model by a neighbor-modulated representation where fitness effects are grouped by recipient of actions (e.g., Hamilton, 1970, Frank, 1998, Rousset, 2004, Fig. 7.1). Second, and as further explained in the main text, one may seek to go one step further and group the indirect fitness effects by actor. We will refer to this as the actor-centered approach to inclusive fitness.

For a model with arbitrary strength of selection on a mutant allele without class structure, a general expression for the decomposition into direct and indirect effects has been reached for the case *n* = 2 by performing a *two-predictor* regression of the fitness of a representative individual from the population, on the mutant allele frequency it carries and on the frequency of the mutant in its neighbors (Queller, 1992; Frank, 1997; Gardner et al., 2011), thus reaching a neighbor-modulated representation of “inclusive fitness”. Importantly, it has been shown that such a *two-predictor* regression automatically yields an actor-centered representation of such effects (Rousset, 2015).

Alternatively, one may perform a single-predictor regression of the individual fitness of a carrier of the mutant on the frequency of the mutant allele among its neighbors, which may be more in line with certain empirical estimates of inclusive fitness where only the social neigborhood of an individual expressing a particular behavior is varied (Krakauer, 2005; Dobson et al., 2012). A single-predictor regression was also used in Lehmann et al. (2016, Box.1) as a justification to derive an exact decomposition of the force of selection into direct and indirect effects for haploid class-structured populations. The single-predictor regression, however, only leads to a neighbor-modulated representation of direct and indirect effects.

In order to obtain a general actor-centered representation of fitness effect for diploid class structured populations and avoid confusions between approaches, we first delineate in the case of haploids the differences between the partitions of fitness by single and two-predictor regressions, and by neighbor-modulated and inclusive fitness effects. In a second time, we turn to the general class-structured populations analysis.

### B.1 Inclusive fitness for haploids without classes

We start by deriving a decomposition into direct and indirect effects from invasion fitness (eq. A.6) for the haploid case and without class structure. To that end, we use the relatedness coefficient defined as

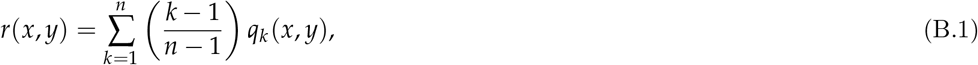

which is the probability that a randomly sampled neighbor of a mutant (itself randomly sampled from its lineage when rare) also carries the mutant allele (when *n* = 2, we have *r*(*x*, *y*) = *q*_2_(*x*, *y*)).

#### B.1.1 Regression with respect to neighbors

For a one-predictor regression, we aim to write the individual fitness of a mutant *x* in a group with trait profile ***x***_*k*_ as

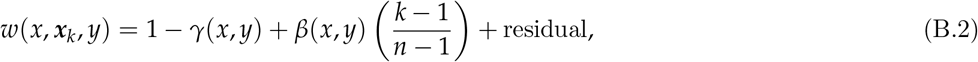

where 1 *− γ*(*x*, *y*) is the intercept of the regression, *β*(*x*, *y*) is the additive effect on a focal’s fitness of allele frequency in neighbors, and (*k −* 1)/(*n −* 1) is the frequency of the mutant allele among neighbors of a mutant. The “cost” (*γ*) and “benefit” (*β*) of this single predictor are determined by minimizing over the *q*_*k*_(*x*, *y*) distribution the expected mean-square difference between individual fitness *w*(*x*, ***x***_*k*_, *y*) and the regression. Thus, for all 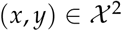, we minimize the sum of squares

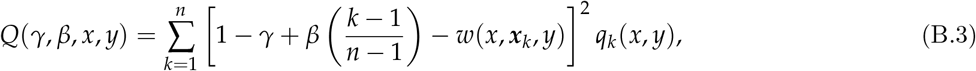

with respect to *γ* and *β*, which are practically obtained by setting *∂Q*(*γ*, *β*, *x*, *y*)/*∂γ*=0 and *∂Q*(*γ*, *β*, *x*, *y*)/*∂β*=0, and solving for *γ* and *β*, which are thus obtained as functions of *x* and *y* (i.e., *γ* = *γ*(*x*, *y*) and *β* = *β*(*x*, *y*)). It follows directly by averaging the regression over the *q*_*k*_(*x*, *y*) distribution, that we can write invasion fitness in terms of the so-obtained coefficient as

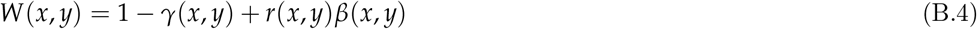

for relatedness defined in eq. (B.1).

#### B.1.2 Regression with respect to focal and neighbors

For the two-predictor regression, the additional predictor variable for the fitness of an individual is its own allelic type. To take this into account in a least-squares regression framework, we need to consider a population where the average mutant frequency is no longer rare. We denote by *p* this frequency, and by a slight abuse of notation, we denote by *w*(*x*, ***x***_*k*_, *p*) the individual fitness of a mutant in a group with a total number *k* of mutant neighbors, in a population where the mutant frequency is *p*. More generally, whenever we will consider fitness at all mutant frequencies, we will replace the last argument of the fitness function with the mutant frequency in the population). Fitness *w*(*y*, ***x***_*k*+1_, *p*) likewise stands for the fitness of an individual carrying the resident allele in the same context of a group including *k* mutants (hence ***x***_*k+1*_ is any vector of dimension *N −* 1 with *k* entries equal to *x* and *N − k* entries equal to *y*). The sum of squares characterizing the regression of the expected number of offspring of a mutant *x* with frequency *p* in a resident *y* population is:

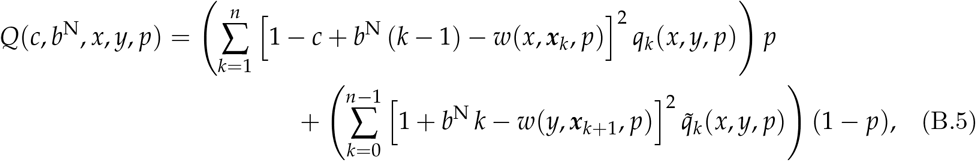

where *c* and *b*^N^ are regression coefficients (the superscript N will from now on stand as a reminder that the regression is, by construction, neighbor-modulated, and we also regress on the number rather than the frequency of mutant neighbors, as this will be useful later), *q*_*k*_(*x*, *y*, *p*) is the probability that, given an individual is a mutant with trait *x* in a population where the frequency of mutants is *p* and residents play trait *y*, it will reside in a group where there are *k* mutants (this probability can be obtained from the full dynamical system for a non-rare mutant, but since we do not need to compute this probability explicitly, we do not specify this dynamical system). Likewise, 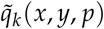 is the probability that, given an individual is a resident with trait *y* in a population where the frequency of mutants with trait *x* is *p*, it will reside in a group with *k* mutants (again this probability can be obtained from the full dynamical system for a non-rare mutant). Minimizing the quadratic form *Q*(*c*, *b*^N^, *x*, *y*, *p*) (by solving *∂Q*(*c*, *b*^N^, *x*, *y*, *p*)/*∂c* = 0 and *∂Q*(*c*, *b*^N^, *x*, *y*, *p*)/*∂b* = 0) we then obtain the regression coefficients *c* = *c*(*x*, *y*, *p*) and *b*^N^ = *b*^N^(*x*, *y*, *p*), which depend on the population allele frequency.

When the mutant is rare (*p →* 0), the fitness of a mutant is *w*(*x*, ***x***_*k*_, *p*) *→ w*(*x*, ***x***_*k*_, *y*) (same as in eq. A.6) and the regression thus predicts this fitness as

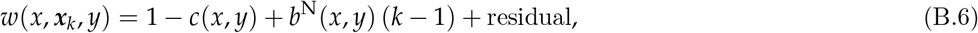

where the residual and the cost and benefit will depend on mutant trait, resident trait, and are limits as *p →* 0 of frequency-dependent terms, i.e., 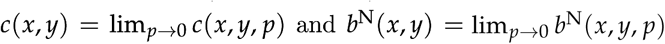. When the mutant is rare, we also have that *q*_*k*_(*x*, *y*, *p*) *→ q*_*k*_(*x*, *y*) because in that case the mutant frequency dynamics within groups is described by the mean matrix ***A***, which is also the matrix of the linearized dynamical system around *p* = 0, and so *q*_*k*_(*x*, *y*, *p*) = *q*_*k*_(*x*, *y*)+ *O*(*p*) for all *k*. The residuals are orthogonal to the regressors when regression coefficients minimize the quadratic form (Cox and Wermuth, 1996, section 3.3.2). Here the regressors include both an intercept and the focal allele frequency, and then the expectation of the residuals is zero whether an individual carries the mutant or the resident allele. Thus, the residuals disappear from the average of expression (B.6) over the conditional distribution *q*_*k*_(*x*, *y*, *p*) of *k* given an individual carries the mutant allele, and we can then write invasion fitness as:

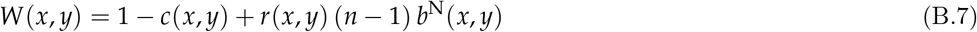

for the relatedness coefficient defined in eq. (B.1). We now give an interpretation of this result. The interpretation of *−c*(*x*, *y*) is the additive marginal effect on the number of successful gene copies produced by an individual when it expresses the mutant instead of the resident allele, while *b*^N^(*x*, *y*) can be interpreted in the two following ways.

1. **neighbor-modulated interpretation.** Here, *b*^N^(*x*, *y*) gives the average effect, on the expected number of offspring (per haplogenome) produced by a focal individual, and stemming from a randomly sampled neighbor expressing a copy of the mutant instead of the resident allele.
2. **Actor-modulated interpretation.** Here, *b*^N^(*x*, *y*) gives the average effect, on the expected number of offspring (per haplogenome) produced by a randomly sampled group neighbor, of an individual expressing a copy of the mutant instead of the resident allele.

This dual interpretation follows from the fact that the regression (e.g. eq. B.5) averages over all contexts, mutant and resident neighbors affecting the fitness of a focal recipient, which itself can be mutant or resident (“two-predictor regression”), and so, on average, recipient and actor individuals can be interchanged (see Rousset, 2015 for more details on this dual perspective). As such, eq. (B.7) provides a genuine actor-centered representation of inclusive fitness and in order to obtain a more compact expression, we let

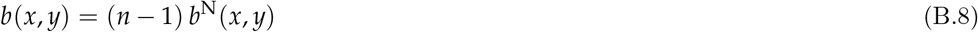

denote the additive effect, on the expected number of offspring (per haplogenome) produced *by all group neighbors*, of an individual expressing a copy of the mutant instead of the resident allele. Thereby, selection favors the mutant (*W*(*x*, *y*) *>* 1) when Hamilton’s rule is satisfied:

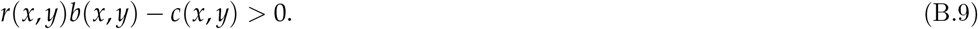

#### B.1.3 Comparing single- and two-predictor regression

The key difference between the single and two-predictor regression version of inclusive fitness (eq. B.3 and eq. B.5) is that only mutant fitness in different contexts (the set of *w*(*x*, ***x***_*k*_, *p*)) are taken into account into the single-predictor regression (eq. B.3), while all contexts for mutant and residents (the set of *w*(*x*, ***x***_*k*_, *p*) and *w*(*y*, ***x***_*k*+1_, *p*) values) are taken into account in the two-predictor version (eq. B.5). Technically, this implies that one has to consider explicitly the average mutant allele frequency *p* in the total population to derive the two-predictor version. Biologically, this implies that the interpretation of costs and benefits differ. Indeed, while the variable *β* in eq. B.4 and *b* in eq. B.7 and are both regression coefficients of fitness to mutant frequency in neighbors, in general *β* ≠ *b*, since the value of a regression coefficient depends on the other predictor variables considered. Likewise, *γ* and *c* differ. This is best seen in the case where *n* = 2, where the single-predictor regression line exactly describes the fitness for *k* = 1 and *k* = 2, hence 1 *− γ* is the fitness of a single mutant in a group. Indeed, in this case

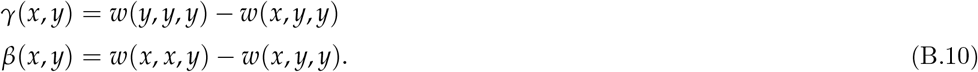

By contrast, in the case of non-additive interactions between group members, it is known that 1 *− c*, as given by the two-predictor regression, is not the fitness of a single mutant (e.g., Gardner et al., 2011, eq. 7). Further, in this case already for *n* = 2, both *c* and *b* will depend on relatedness coefficients (e.g., Gardner et al., 2011) and are given explicitly by

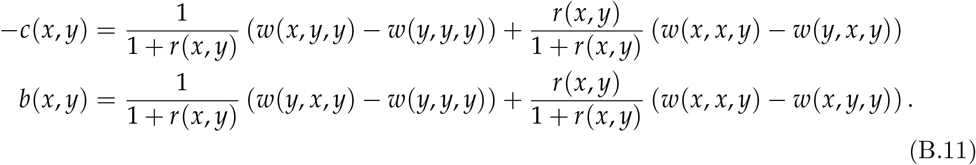

In the actor-modulated interpretation, 1/[1 + *r*(*x*, *y*)] and *r*(*x*, *y*)/[1 + *r*(*x*, *y*)] weigh in both *−c* and *b* the case where the neighbor of a focal individual is either resident or mutant, respectively. To understand where these weights come from, let *P* denote the probability of an (*x*, *x*) focal-neighbour pair in a group and *Q* the probability of an (*x*, *y*) focal-neighbour pair [these probabilities reducing respectively to *pr*(*x*, *y*) and to *p*(1 *− r*(*x*, *y*)) for vanishing *p*]. Then the weights are proportional to *P* + *Q* versus *P*, rather than *Q* versus *P*, for the following reason. The derivative of the sum of squares with respect to *c* is proportional to

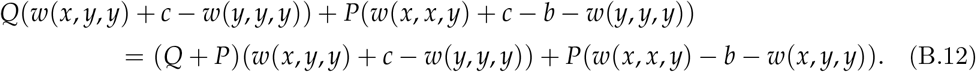

The effect of least-square regression is to predict *w*(*x*, *y*, *y*) and *w*(*y*, *x*, *y*) with identical prediction residuals: *w*(*y*, *x*, *y*) *− w*(*y*, *y*, *y*) *− b* = *w*(*x*, *y*, *y*) *− w*(*y*, *y*, *y*) + *c*, and thus the derivative is proportional to

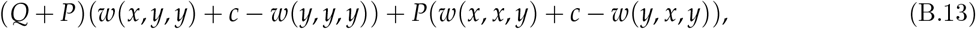

meaning that we have represented the original term *w*(*x*, *x*, *y*) *− w*(*y*, *y*, *y*) as the effect of two allelic substitutions, each with effect *−c*. This recovers the solution for *−c*. The same logic holds for the weights in the expression for *b* and this argument holds at all allele frequencies *p*.

### B.2 Inclusive fitness for diploids with classes

#### B.2.1 Multiplayer class-structured regression

We now turn to deriving an expression for inclusive fitness for diploids with class structure by performing an extension of the two-predictor regression of the fitness of a representative gene copy from the population. To construct this fitness measure, we consider all possible group trait profiles, and write for each class *u* of descendants and each class *s* of parent, the fitness (per haplogenome) of a focal individual *i* with trait *x*_*i*_ in a group with trait profile ***x***_−*i*_ as

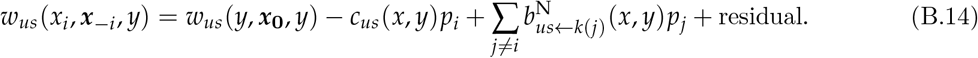

Here, *p*_*i*_ denotes the frequency of the mutant allele in individual 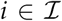 [zero, one-half, or one, so that the individual expresses, respectively, trait *y*, *x* or *z*(*x*, *y*)]. The function 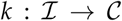 assigns to each individual in group 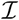 its class in 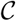, such that *k*(*j*) = *a* when individual *j* is in class *a*. The mutant allele frequencies *p*_*i*_ in each individual are thus predictor variables of fitness and *c*_*us*_(*x*, *y*) and 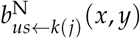 are regression coefficients depending on mutant and resident trait values. Indeed, eq. (B.14) says that we seek to obtain the predictor

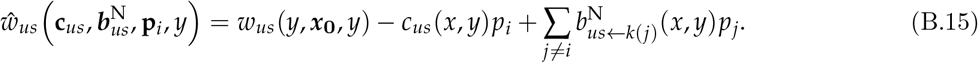

of class-*u* fitness of a class-*s* gene copy as a linear regression on the mutant allele frequency carried by all actors on that fitness [here the vector 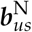 collects all the *b*_*us←k*(*j*)_ regressors and the vector **p**_*i*_ collects mutant frequencies *p*_*i*_ in all individuals].

Eq. (B.14) must hold for all trait profiles 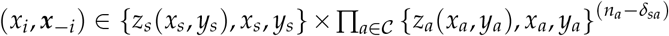 and the regression coefficients are determined by the following argument, which generalizes the one developed in the absence of class structure (section B.1). Let then *w*_*us*_(*x*_*i*_, ***x**_−i_*, ***p***) denote the fitness of an individual in a population where the mutant frequencies in the different classes are no longer rare, and are collected in the vector 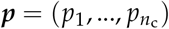, where *p*_*s*_ is the average mutant allele frequency in class *s* in the population. Further, let 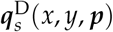 denote the ordered distribution of group traits, determined by the distribution of contexts of copies of the mutant allele in class *s*. This 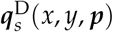 distribution has the same sample space as 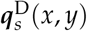) (recall eq. A.30) and generalizes it to arbitrary allele frequency. Likewise, 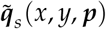 denotes the ordered distribution of group traits for non-rare mutant frequency, determined by the distribution of contexts of copies of the resident allele in class 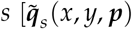 has sample space 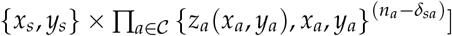. With these notations, the expected sum of squares to be minimized by the regression coefficients can be written

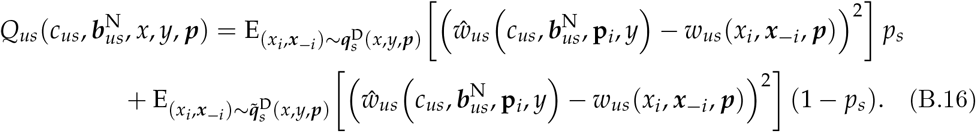

By solving 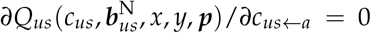 and 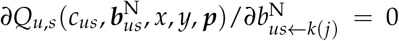 for all *j* ≠ *i*, we obtain the regression coefficients *c*_*us←a*_(*x*, *y*, ***p***) and 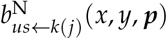. These depend on the population state and we note that that

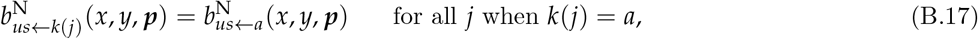

since individuals from the same class carrying similar traits have the same effect on the recipients of their actions. Computing the regression coefficients explicitly involves solving linear systems of equations with complicated terms. But all in all, this is not more involved that computing invasion fitness to begin with, since this requires computing eigenvectors (recall eq. A.8). Hence, no computational complexity is added if the regression coefficients are to be computed explicitly. That said, the interpretative nature of inclusive fitness does require the explicit computation of the regression coefficients.

#### B.2.2 Average allele frequency

Our aim is now to evaluate the so-obtained regression coefficients under vanishing mutant allele frequency. To do this, we need a single (scalar) measure of allele frequency such that allele frequencies in all classes vanish simultaneously when this measure vanishes. As such a measure, we use the weighted average allele frequency 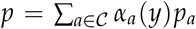 in the population, where the weights are the neutral class reproductive values (the *α*_*a*_(*y*) = *v*_*a*_(*y*)*ϕ*_*a*_(*y*, *y*) elements in eq. A.26). To evaluate the regression coefficients, we then need to be able to express each class-specific frequency *p*_*a*_ in terms of *p* and *ϕ*_*a*_(*x*, *y*), at least when the mutant allele is rare. For this purpose, we recall that as long as the mutant allele is rare, its growth is characterized by the leading eigenvalue (invasion fitness) and by the associated right eigenvector (quasi-stationary distribution) ***u***(*x*, *y*) of the transition matrix ***A***(*x*, *y*) [i.e., eq. A.8]. Eigenvectors are defined up to a constant factor, so the relationship between allele frequencies *p*_*a*_ in each class *a* and the eigenvector can be specified up to a constant, here denoted *L*_1_. We write this relationship as

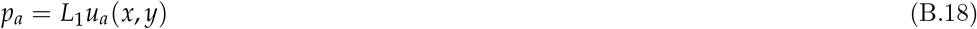

where *u*_*a*_(*x*, *y*) is (up to a constant factor) the frequency of the mutant allele in class *a* under the quasi-stationary distribution ***u***(*x*, *y*). The average allele frequency is then 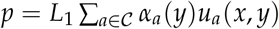, whereby 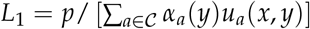 and

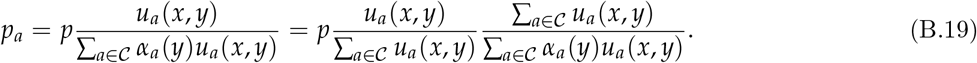

From eq. (A.22), the middle fraction on the right-hand side is the probability *ϕ*_*a*_(*x*, *y*) that a randomly sampled gene copy from the mutant lineage is in class *a*, introduced in eq. (A.21): 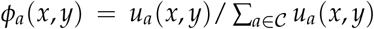. The last fraction in eq. (B.19) is then the inverse of the fraction 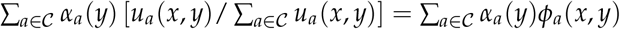, and

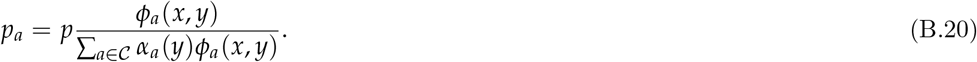

Substituting eq. (B.20) into *c*_*us*_(*x*, *y*, ***p***) and 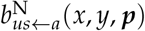, and recalling eq. (B.17), we compute the regression coefficients of eq. (B.14) as

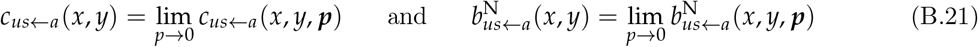

(see section B.3 for a concrete application and explicit computation of such coefficients). We further note that, by construction, ***q***_*s*_ (*x*, *y*, ***p***) *→ **q***_*s*_ (*x*, *y*) as *p* → 0. This then allows us to define

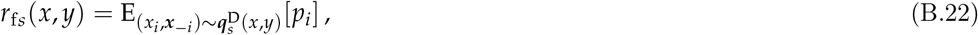

which is the probability that, conditional on a random gene copy of class *s* carrying the mutant allele, a randomly sampled homologous gene in that individual is a mutant. For *k*(*j*) = *a*, we have

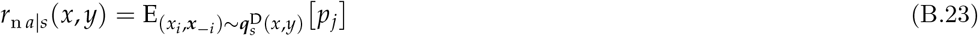

which is the probability that, conditional on a random gene copy of class *s* carrying the mutant allele, a randomly sampled homologous gene in a neighbor of class *a* is a mutant allele. In terms of the *r*_f*s*_(*x*, *y*) and *r*_n*a|s*_(*x*, *y*) probabilities, we define the relatedness coefficient between a class-*s* actor and a class-*a* recipient as

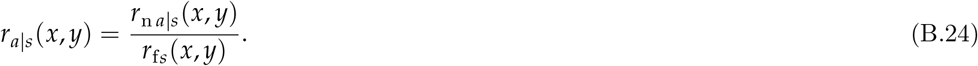

#### B.2.3 Average inclusive fitness

Now substitute eq. (B.14) into direct fitness (eq. A.29) and then into invasion fitness (eq. A.27). Then, by dint of the reproductive values recursion (eq. A.18), the reproductive values normalizer (eq. A.20), the relatedness coefficients (eqs. B.22–B.24), the relationship 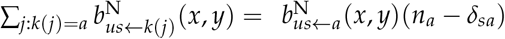 (eq. B.17), and recalling that the residual term in eq. (B.14) cancels when averaged over the 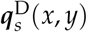 distribution (since they are uncorrelated with regressors), the invasion fitness of a mutant allele introduced as a single copy in a resident population can be put under the form

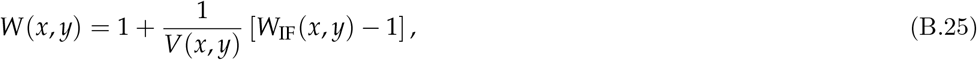

where

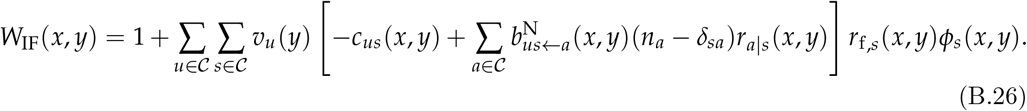

is the inclusive fitness of the mutant allele. These two equations were previously derived for the haploid case (*r*_f,*s*_(*x*, *y*) = 1 for all 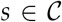) in Lehmann et al. (2016, eqs. C.1–C.6) assuming that 1 *− c*_*u,s*_ was the intercept and only 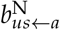 were the regression coefficients of fitness, thus performing a multiple-neighbors extension of the single-predictor regression to obtain inclusive fitness. To obtain such coefficients it suffices to set *p*_*s*_ = 1 in eq. (B.16) and otherwise follow the same line of argument.

Since *V*(*x*, *y*) *>* 0, we have from eq. (B.25) that

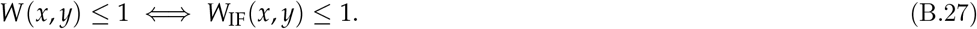

Hence, trait *x*^∗^ is uninvadable if it is a best-reply to itself in terms of inclusive fitness: 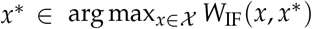. Finally, we note that if one chooses to normalize the (neutral) reproductive such that *V*(*x*, *y*) = 1, then one would have *W*(*x*, *y*) = *W*_IF_(*x*, *y*).

#### B.2.4 Grouping effects by actor

In eq. (B.26), the fitness effects of social interactions among individuals in the population are grouped by recipients each class *s*. We now first rearrange eq. (B.26) in order to obtain a grouping of fitness effects by actor in each class, so that *W*_IF_(*x*, *y*) reads as an average over class-specific inclusive fitnesses (which justifies the subscript “Inclusive Fitness”). In a second, time we then show that class-specific inclusive fitness is maximized in an uninvadable population state.

The key steps to reach the actor-centered perspective, is to note, first, that, as was the case under the haploid model, 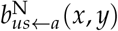 can be interpreted in the two following ways.

1. **Neighbor-modulated interpretation.** Here, 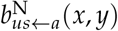 gives the average effect, on the expected number of class-*u* offspring (per haplogenome) produced by a focal individual in class *a*, and stemming from a randomly sampled neighbor expressing a copy of the mutant instead of the resident allele.
2. **Actor-modulated interpretation.** Here, 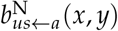 gives the average effect, on the expected number of class-*u* offspring (per haplogenome) produced by a randomly sampled group neighbor of class *a*, of an individual expressing a copy of the mutant instead of the resident allele.

We will thus from now on use the second interpretation and further note that the following equality holds

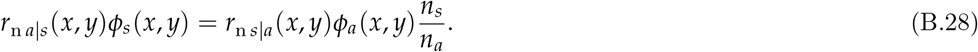

To check this result, we highlight that each side of the equation involves two ways of sampling gene copies. First, we sample gene copies uniformly from the mutant lineage (by definition, *ϕ*_*s*_(*x*, *y*) is the probability that a gene sampled in this way is in a class-*s* individual), and then we sample gene copies uniformly among class-*a* individuals (*r*_n*a|s*_(*x*, *y*) is the probability that, when a given gene copy from a class-*s* individual is mutant, a given gene copy from a class-*a* individual in the same group is mutant). The expected number of pairs of gene copies in class-*s* and class-*a* individuals within a group per copy of the mutant allele is then obtained as 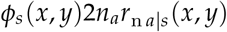 when the first sampled gene copy is in a class-*s* individual, and as 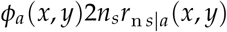 when the first copy is sampled in a class-*a* individual, from which the above result follows.

Substituting eq. (B.28) into eq. (B.26), and rearranging we obtain

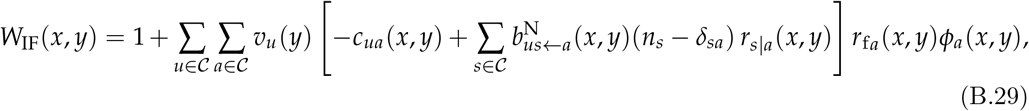

where

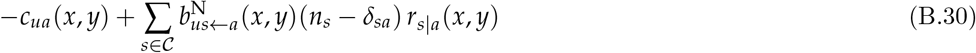

is the average effect, on the number of class-*u* offspring (per haplogenome) produced by all group members of class *a*, and stemming from an individual of class *a* expressing a copy of the mutant instead of the resident allele. Eq. (B.30) is consistent with eq. (8) of Grafen, 2006a who assumed (a) additive separable fitness effects and (b) relatedness independent of evolving trait values. To further simplify expression (B.29), we let

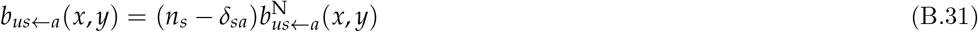

denote the average additive effect, on the number of class-*u* offspring produced (per haplogenome) by all class-*s* neighbors in a group, and stemming from a single class-*a* individual switching to expressing a copy of the mutant instead of the resident allele. Substituting eq. (B.31) into eq. (B.29), we can obtain:

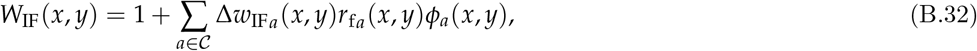

where

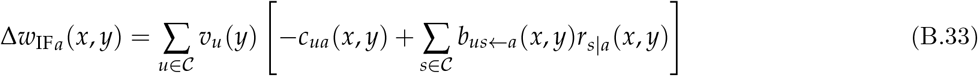

is the inclusive fitness effect of an average class-*a* carrier of the mutant allele.

#### B.2.5 Class-specific inclusive fitness maximization

We here prove that the inclusive fitness effect (eq. B.33) is maximized with respect to *x*_*a*_ at the uninvadable state *x*^∗^, which will allow us to define a class-specific inclusive fitness that is equivalently maximized. In general, inclusive-fitness (eq. B.32) maximization does not implies maximization of the summand therein for each class with respect to all mutants 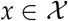, but what we consider here are class-specific mutants. To that end, let 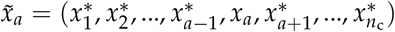, denote the trait profile of a mutation with all traits held at the uninvadable state, except for trait *x*_*a*_ of class *a* that can unilaterally deviate. Then, if the mutant 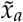 appears in a population in state *x*^∗^, we have

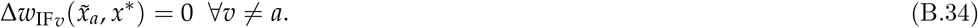

Eq. (B.34) says that if a mutant allele changes only the trait expression of individuals of class *a*, then the inclusive fitness effect of any other class *v* ≠ *a* is nil. This is so since 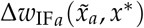 captures all effects of individuals of class *a* expressing the mutant trait *x*_*a*_ (the “actors”) on mutant allele transmission. A more formal proof follows from the fact that 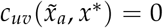 and 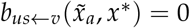 for all *v* ≠ *a* and all *u* and *s* because the *u*-type fitness *w*_*us*_ of an individual of class *s* is a constant with respect to the traits of individuals in any class *v* ≠ *a*, since all individuals in any class *v* ≠ *a* express the same trait value 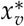. Hence, all such regression coefficients on class-*v* individuals will be nil, since there is no variation in individual fitness to be explained by any such regressor.

Substituting eq. (B.34) into eq. (B.32), the inclusive fitness of a mutant inducing an unilateral deviation in class *a* in population state *x*^∗^ is given by

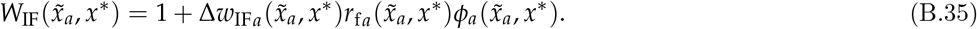

Since 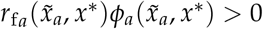 for all classes and traits, we have from eq. (B.35) that

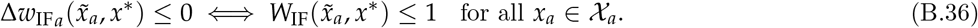

Hence, trait 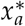 preempts invasion by any mutant inducing an unilateral deviation in class *a* if it satisfies 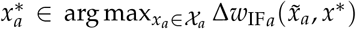. Since this holds for each class in an uninvadable population *x*^∗^, we have

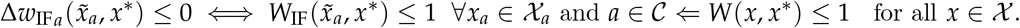

In other words, the inclusive fitness effect in each class is maximized in an uninvadable population state:

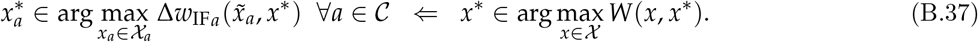

The left-hand side can now also be written in terms of the inclusive fitness

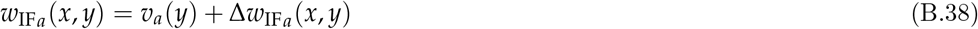

of a class-*a* individual (eq. 4 of the main text), which is thus also maximized in an uninvadable population state (since *v*_*a*_(*y*) does not depend on the mutant trait):

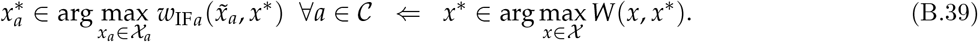

We finally mention that if we were to replace class-specific inclusive 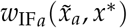 by classspecific average direct fitness 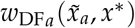 in the left-hand side of eq. (B.39), then eq. (B.39) is not satisfied, unless additional assumptions are made on the form of the individual fitness functions. This negative results points to the limitations of using average direct fitness as an individual-centered maximand. Indeed, in the presence of indirect fitness effects, where an actor of class *a* carrying the mutant allele affects the fitness of another individual carrying the mutant (say a worker affecting the reproduction of a queen), the direct fitnesses 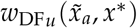 for *u* ≠ *a* in expression (A.27) for invasion fitness will not be independent of the mutant trait *x*_*a*_ of a class-*a* individual. In this case, the invasion fitness of trait *x*_*a*_ depends on these 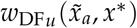 fitnesses for *u* ≠ *a*, even though the individual expressing trait *x*_*a*_ is not in any class *u* ≠ *a*. Hence the biological interpretation of an individual of class *a* as maximizing 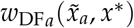 at an evolutionary equilibrium generally breaks down.

### B.3 Example: inclusive fitness for social insects

We here derive the inclusive fitness effects for the social insect model (eq. B.5 in Box 2) from the fitness functions defined in the main text (see eqs. B.2–B.4 in Box 1) and assuming the population has reached the uninvadable sex ratio of 1/2 for this model. Because we consider only diploidy, our “social insects” are akin to termites rather than ants. From eqs. (B.2) of Box 2, we can write the individual fitness of a female *i* whose worker offspring has trait *x*_o(*i*)_ ∈ {*z*_o_, *x*_o_, *y*_o_} as

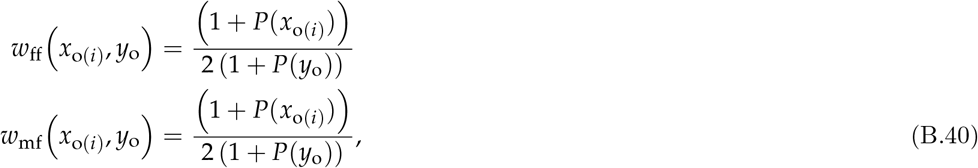

which, once averaged over the cases where the offspring is heterozygote (with phenotype *x*_o(*i*)_ = *x*_o_), or homozygote resident (with phenotype *x*_o(*i*)_ = *y*_o_), produces eqs. (B.2) of Box 2 (and where for simplicity of presentation we only denote the traits whose variation affect fitness). Likewise, from eq. (B.4) of Box 2, the individual fitness of a male *i* whose worker offspring has trait *x*_o(*i*)_ ∈ {*y*_o_, *x*_o_, *z*_o_} can be written

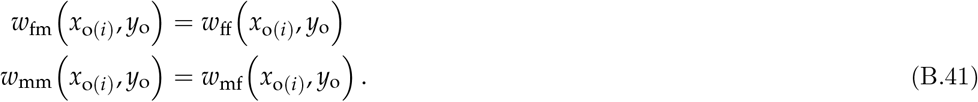

In order to evaluate the sum of squares for the regression coefficients, we need to take into account all possible matings as this determines the number of mutant allele copies in the worker offspring. There is a total number of 9 matings, since a female can be homozygote mutant (probability denoted *p*_ho,f_), heterozygote (probability denoted *p*_he,f_), or homozygote resident (probability (1 *− p*_ho,f_ *− p*_he,f_)), and her mate can be of the same respective types (with respective probabilities, *p*_ho,m_, *p*_he,m_, and (1 *− p*_ho,m_ *− p*_he,m_)). The assumption that we consider a population with random mating at the uninvadable sex-ratio, implies that the fitness functions for both males and females are equivalent (e.g., eq. B.41), and that the frequency of the mutant allele will be the same in males and females, *p*_m_ = *p*_f_ = *p*. Henceforth, we can evaluate the genotype frequencies in terms of allele frequencies at Hardy-Weinberg equlibrium:

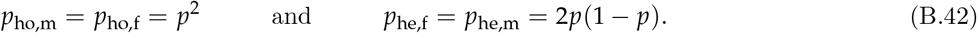

#### B.3.1 Regressions for female fitness components

Taking into account all matings, we write the sum of squares for female fitness through offspring of type *j* ∈ {*f, m*} as

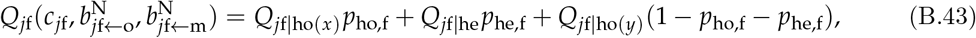

where *Q*_*jf|ho*(*x*)_, *Q*_*jf|he*_, and *Q*_*jf|ho*(*y*)_ are, respectively, the sum of squares when the female is homozygote mutant, heterozygote, and homozygote resident. Application of eqs. (B.15)–(B.16) shows that when the female is homozygote

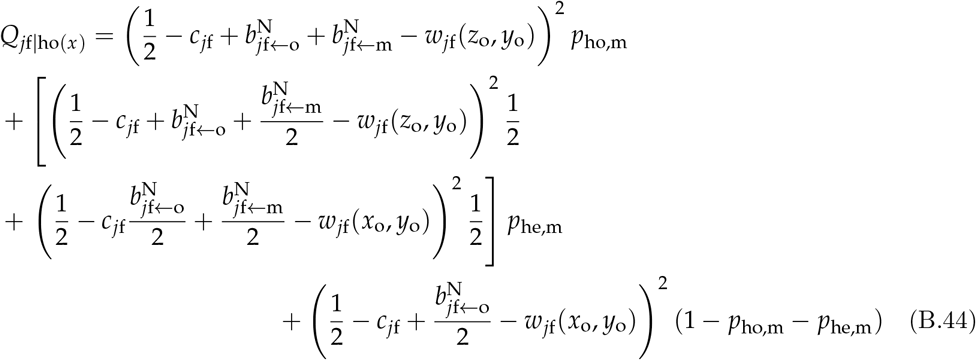

where in the present example *w*_*jf*_ is given by eq. (B.40). The first, second, and third summand, stand, respectively, for the case where the male mate of the focal female is homozygote mutant, heterozygote, or homozygote resident. When the male is heterozygote, then with probability 1/2 the worker inherits a copy of his mutant allele and will be homozygote (first term in the second summand), while with probability 1/2 the worker does not inherit a copy of the mutant allele from its father and will be heterozygote (second term in the second summand).

When the female is heterozygote, we write the sum of squares as *Q*_*jf|he*_ = (1/2)*Q*_*jf|he*,1_ + (1/2)*Q*_*jf|he*,0_, where *Q*_*jf|he*,1_ represents the case where the worker inherits the mutant allele from its mother and *Q*_*jf|he*,0_ for the case the worker does not inherit the mutant from its mother. We find that

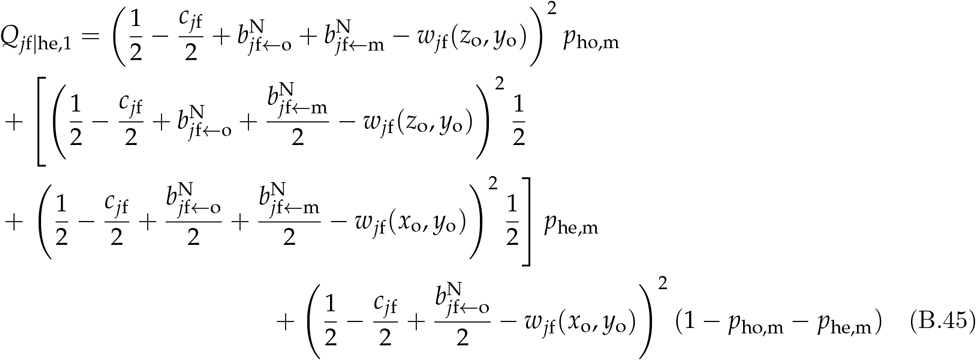

and

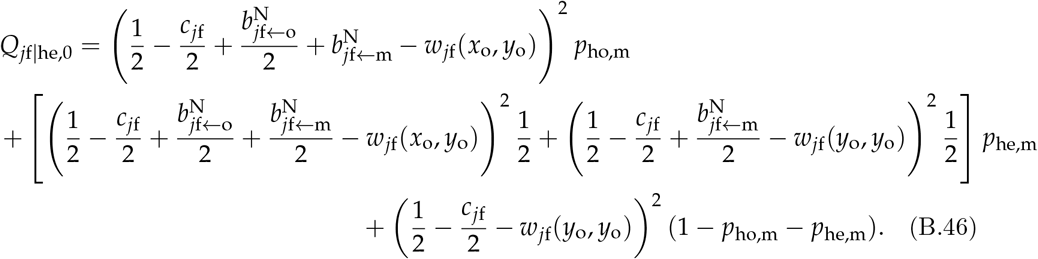

Finally, when the female is homozygote resident, we have that

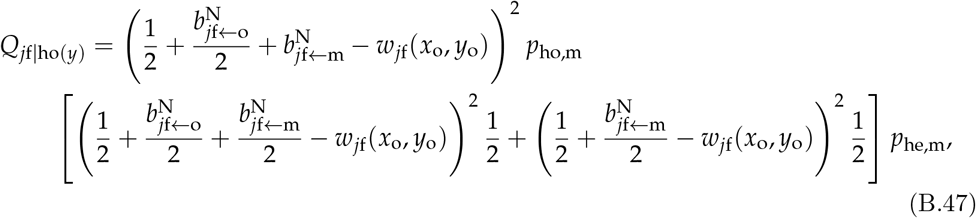

since a worker from a homozygote resident mother can inherit the mutant allele only from its father, and when the father is heterozygote the worker inherits the mutant with probability 1/2.

We now minimize the sum of squares 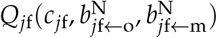 with respect to the relevant regression coefficients, which requires that, for *j* ∈ {*f, m*}, we solve

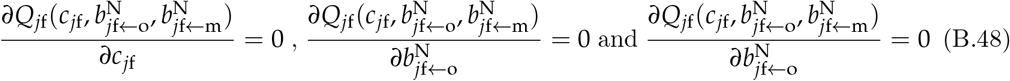

for *c*_*jf*_, 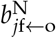 and 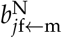. Substituting eq. (B.42) into the so-obtained regression coefficients and letting *p →* 0, we finally obtain that *c*_*jf*_ = 0, 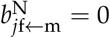 for *j* ∈ {*f, m*}, and

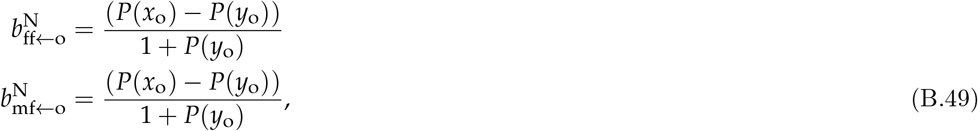

where these regressions coefficients are the offspring production of the mutant worker minus that of the resident worker relative to the offspring production of the mutant worker.

#### B.3.2 Regressions for male fitness components

We now derive the regression coefficients for the male fitness components. The model for the male side is exactly symmetric to that of the female side and to compute the corresponding sum of squares 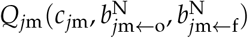 for *j* ∈ {*f, m*} we only interchange m and f subscripts in all equations of the previous section. Otherwise, the calculations carry over *mutatis mutandis* to give *c*_*jm*_ = 0, 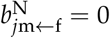 for *j* ∈ {*f, m*}, and

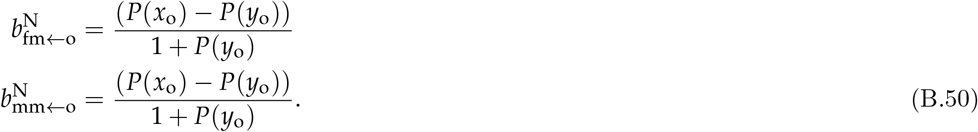

#### B.3.3 Inclusive fitness effects

Using the regression coefficients computed in the last two sections, we are now in the position to compute the inclusive fitness effects. First, the inclusive fitness effects of females and males is null

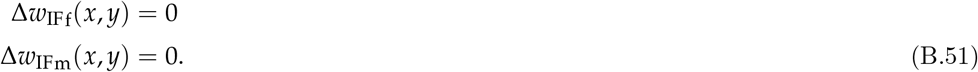

To obtain the inclusive fitness effect for a worker, we note that from eq. B.31,

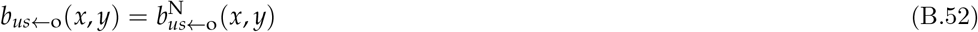

for *u ∈ {*f, m*}* and *s ∈ {*f, m*}* since the number of individuals of each class *n*_f_ = *n*_m_ = *n*_o_ = 1. With this, eq. (B.33), and eqs. (B.49)–(B.50), we obtain

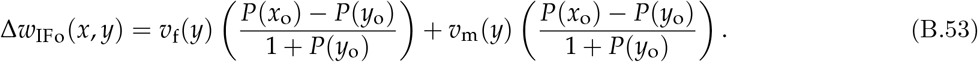

Adding the reproductive values to the inclusive fitness effects shown in eq. (B.51) and eq. (B.53), we obtain the inclusive fitnesses displayed in Box 2.

## Supplement C: Fitness as-if

### C.1 Rational-actor payoff maximization

As explained in the section “Fitness as-if” of the main text, a standard concept for the prediction of individual behavior is that of a Nash equilibrium trait profile, compared to which no individual can get a higher “payoff” by a unilateral deviation of behavior (see e.g., Luce and Raiffa, 1957, Fudenberg and Tirole, 1991 or Mas-Colell et al., 1995). Let us now introduce such a payoff function for a class *a* individual, denoted *w*_I*a*_, and defined on the domain

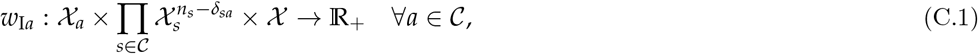

such that 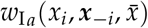 is the payoff to individual *i*, 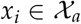 being now understood as any trait value that a given individual *i* could express instead of the trait that it actually expresses, when group neighbors express trait profile ***x**_−i_*, and in a population where the average trait expression is 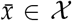. This payoff function has exactly the same domain as the individual fitness function (eq. A.3).

Suppose now each individual in the population is envisioned as an autonomous decision-maker, “choosing” the trait it expresses independently of each other individual and with a striving to maximize its payoff function *w*_I*a*_ (hence *x*_*i*_ can vary and is not genetically determined). Then, a Nash equilibrium 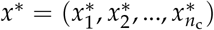 (symmetric in each class) satisfies

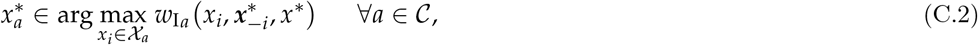

where 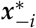 is the trait profile of all neighbors, where entry *j* of 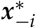 is equal to 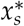 if neighbor *j* ≠ *i* is of class *s*. In such a symmetric Nash equilibrium 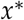, individuals in each class make the best decision for themselves in terms of payoff, i.e., they maximize their payoff based on all others doing the same.

Our aim is to elicit a representation of the payoff function *w*_I*a*_ that individuals appear to maximize in an uninvadable population state. We call such a payoff a fitness as-if. More formally, a fitness as-if function *w*_I*a*_ satisfies

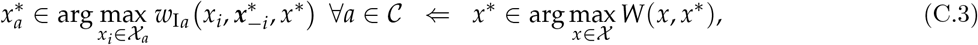

where the invasion fitness in the right-hand side is given by eqs. (A.27)–(A.29). Eq. (C.3) says that if *x*^∗^ is uninvadable, then this equilibrium can be envisioned as a Nash equilibrium, where each individual appears to maximize its fitness as-if, when each other individual in each class exhibits fitness-maximizing behavior. In other words, in an uninvadable population state, it is as if each individual maximizes its fitness as-if.

In this Supplement, we not only prove that eq. (6) of the main text satisfies eq. (C.3), but more generally explain how to construct expressions for fitness as-if that take the form of both average direct fitness and inclusive fitness. Thereby, this Supplement fully connects together traditional game theory, evolutionary invasion analysis and inclusive fitness theory.

### C.2 Average direct fitness as-if

#### C.2.1 The instrumental distribution

We start by presenting a way to construct an average direct fitness as-if as this will pave the way to construct inclusive fitness as-if. Since we aim that the trait of each individual and thus of neighbors can be distinct from each other, we need to depart from the population genetic models of the previous sections where invasion fitness was depending only on heterozygote and homozygote mutant and resident traits, with the distribution ***q***^D^(*x*, *y*) of group states describing correlated trait expression within groups. In order to take this difference into account, we consider that, while fitness as-if should in general consist of the same fitness components, *w*_*us*_, as invasion fitness, it should be averaged over a different distribution of correlated trait expression within groups (in particular, a distribution with a different sample space allowing for each individual expressing a different trait). We refer to this new distribution as the *instrumental* distribution, and it will be reminiscent of the so-called subjective probability distribution of the profile of traits that neighbors play, as considered in the construction of an individual’s utility function in game theory (e.g., Fudenberg and Tirole, 1991, Mas-Colell et al., 1995). To describe how we obtain the instrumental distribution, we first define its sample space, beginning with a haploid population without class structure.

#### C.2.2 Haploids without classes

For haploids without classes, where 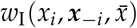 is the fitness as-if of an individual with trait *x*_*i*_ in a group with neighbor trait profile ***x***_−*i*_ = (*x*_1_, *x*_2_, …, *x*_*i−1*_, *x*_*i+1*_, …, *x*_*n*_) in a population with average group trait 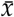, the instrumental distribution is constructed as follows. We first consider the sample space defined from the neighbor trait profile ***x***_−*i*_, defined by replacing any number of the elements of ***x***_−*i*_ by *i*’s trait. Thus, for any *k ∈ {*1, …, *n*}, we consider the set 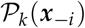 of hypothetical neighbor trait profiles 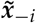 such that exactly *k −* 1 components of the profile ***x***_−*i*_ are replaced by *i*’s trait *x*_*i*_, while the remaining *n − k* components of 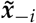 are identical to those in ***x***_−*i*_ (this operation will capture correlated trait expression within groups). The set of all such profiles is 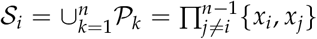. From the perspective of individual *i*, we can think of 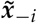 as a hypothetical profile where neighbors’ traits have been replaced with traits similar to self, and if such a profile were to obtain in individual *i*’s group, then its fitness would be 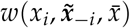).

Any probability distribution 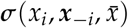 on the space 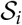 takes values in the simplex 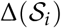 induced by 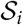, and assigns probabilities 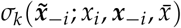 such that these probabilities satisfy

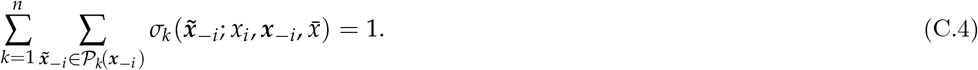

The instrumental distribution 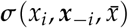 is as yet undefined beyond its sample space. In particular, this distribution has yet no imposed relation to the probabilities of events specified by the ***q***^D^(*x*, *y*) distribution that occur in the actual reproductive process in the population under consideration, but it retains the ability to describe within-group correlated trait expression. In order to connect these two distributions, we note that ***q***^D^(*x*, *y*) takes values in the simplex 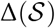 or ordered phenotypic groups states, which is the same as the simplex 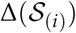. Thus, ∆(*S*) = 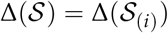, and we can choose the instrumental distribution such that

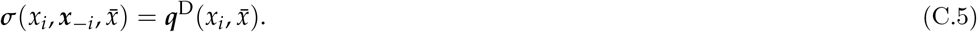

So we consider that the instrumental probabilities of events, defined by replacing elements of the trait profile by the focal individual’s trait, are identical to the probabilities of ordered trait profiles in the population genetic model (or equivalently, to the probabilities of joint genetic identity in the group).

Given the instrumental distribution defined by eq. (C.5), we can define the average direct fitness as-if of an individual with trait *x*_*i*_ as

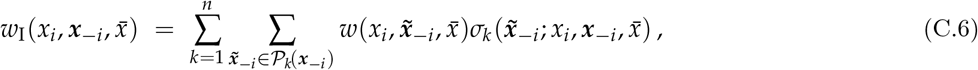

which is the average of individual fitness over the distribution 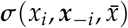. A more compact representation of this fitness as-if is

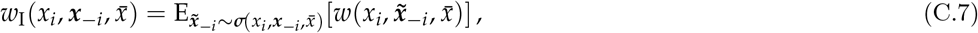

where the notation *∼* specifies that variable 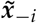 follows the distribution 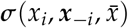 (recall eq. A.12).

#### C.2.3 Diploids with classes

We can now generalize the construction of the instrumental distribution to a diploid class-structured population. For this case, it is useful to denote explicitly by *x*_*i,a*_ the realized trait of individual 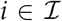 when of class *a*. Then, the instrumental distributions 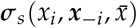 for the realized profile of traits 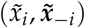 in a group when individual *i* is of class *a* is defined as follows. First, the trait 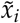 of individual *i* when of class *a* is assumed to take values in 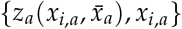 and this parallels the case where in the genetic process an individual with the mutant allele can be heterozygote or homozygote, but our assignment is here a defining feature and is not intended to reflect any genetic reality. Second, each element 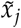 of 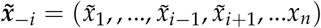 is defined to take values in the set of traits belonging to the class of the individual under scrutiny; that is, if individual *j* is of class *s* then, by construction 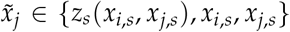. The hypothetical profile 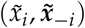 is then defined to be distributed according to 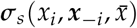, a distribution that has sample space

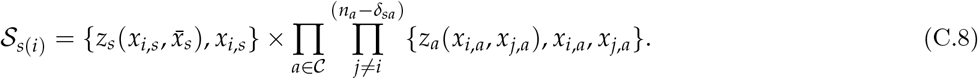

Hence, the distribution ***σ**_s_* takes values in the set 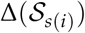, which is the simplex generated by the support 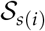. Since the distribution 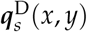 determining invasion fitness takes values in the same simplex 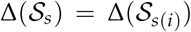 (recall eq. A.30), we can, as in the haploid case, set the probabilities of events in the instrumental distribution identical to the probabilities of ordered trait profiles in the population genetic model:

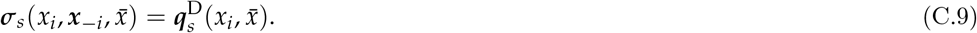

Thus, we again define the instrumental probabilities, of events defined by replacing elements of the trait profile by the focal individual’s trait, to be identical to the probabilities of ordered trait profiles in the population genetic model.

Given the instrumental distribution 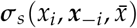 so defined (eq. C.9), the average direct fitness as-if of an individual of class *a* with trait *x*_*i*_ in a group with neighbor trait profile ***x**_−i_* in a population with average group trait 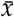 is defined as a reproductive value-weighted sum of expected numbers of offspring of different classes *u*:

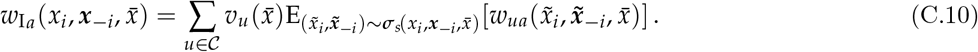

In order to illustrate the notation and better understand the expectation in eq. (C.10) for diploidy, we consider the case of *n* = 2 without class structure (hence the neighbor trait profile is the singleton ***x***_−*i*_ = *x_−i_*). Then, we write the direct fitness as-if of an individual with trait *x*_*i*_ as

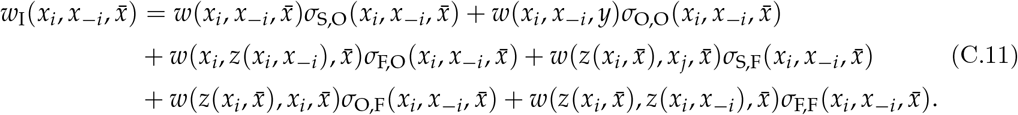

Here, the second subscript *k ∈ {*O, F} in 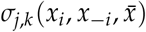 denotes that the instrumental substitute to individual *i* can be of two possible types, either it is “outbred” (*k* = O), in which case its (objective) fitness *w* depends on trait *x*_*i*_, or it is “inbred” (*k* = F), in which case its fitness depends on trait 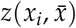. The first subscript *j ∈ {*S, O, F} denotes that the instrumental substitute to the group neighbor can express three different traits: it expresses either trait *x_−i_* (*j* = S for “self”), or *x*_*i*_ (*j* = O), or 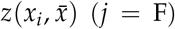. With these notations, 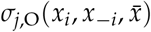 is the instrumental probability that, given trait profile 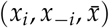, individuals *i* is of type “outbred” and its neighbor expresses the trait of type *j ∈ {*S, O, F}, while 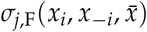 is the instrumental probability that individual *i* is “inbred” and its neighbor expresses trait of type *j*. In terms of these probabilities, we can write the instrumental distribution of profiles experienced by individual *i* as

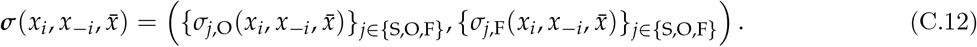

Note that eq. (C.11) shows that fitness as-if is defined as an average over cases where individuals are “outbred” or “inbred”, i.e., have the trait of an heterozygote or homozygote, and so varying *x*_*i*_ varies the trait both when the substituted individual is heterozygote (given by *x*_*i*_ itself) and when it is homozygote (given by *z*(*x*_*i*_, *x*_*j*_)). This construction ultimately owes to the fact that in the original reproductive process individuals express different traits upon being heterozygote or homozygote (e.g., eq. A.29), a standard modeling assumption for diploids (e.g., Nagylaki, 1992; Gillespie, 2004; Hartl and Clark, 2007).

### C.3 Inclusive fitness as-if

#### C.3.1 Multiplayer regression for class-structure

We now present the regression analysis underlying the construction of inclusive fitness as-if and do so directly for diploids with class structure. To that end, we consider the same regression model as in the population genetic model (recall eqs. B.14–B.15) but will evaluate its coefficients under the instrumental distribution instead of the genetic contextual distribution. Namely, we focus on individual *i* with trait *x*_*i*_ in a group with neighbor trait profile ***x**_−i_*, and consider a hypothetical switch in behavior to expressing trait 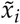 in a group with neighbor trait profile 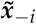. We then write the number *u* of descendants of an individual *i* of class *s* in the altered group as

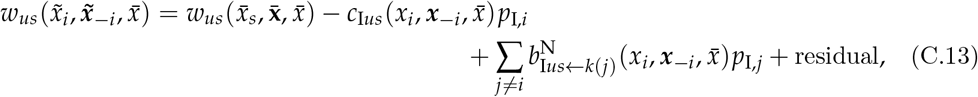

where 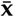 denotes the vector of neighbor trait profile (dimension *n −* 1), all of which are evaluated at the mean (of the corresponding class) in the population and 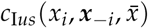 and 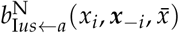 are regression coefficients. The functional form of this equation is the same as that under the population genetic model (eq. B.14), but its interpretation slightly differs and is as follows. We consider a group state where each of the gene copies from the original group of the focal individual *i* may be replaced in any individual by 2, 1, or 0 copies of an I allele, which plays the same role as the mutant allele in being a determinant of the traits of a focal individual *i* and its neighbors. For the focal individual itself, the new trait value 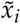 is within the set 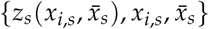, when it bears 2, 1, or 0 copies of the I allele, and *p*_I,*i*_ ∈ {0, 1/2, 1} denotes the frequency of allele I in individual *i*. The new trait 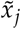 of a neighbor individual *j* ≠ *i* when of realized class *k*(*j*) = *a* is within the set {*z*_*a*_(*x*_*i,a*_, *x*_*j,a*_), *x*_*i,a*_, *x_j,a_*}, which stems, respectively, from individual *j* ≠ *i* bearing 2, 1, or 0 copies of the I allele. Thus any value of 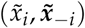 is a hypothetical group trait profile, resulting from a switch of allele expression and where, by construction, eq. (C.13) must hold for all profiles 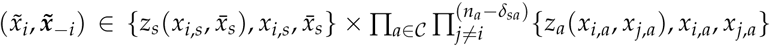. Hence, the main structural difference between eq. (C.13) and its population genetic counterpart, eq. (B.14), is that all neighbors of individual *i* have all distinct traits in eq. (C.13) when they do not switch to expressing allele I.

The regression coefficients 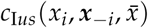 and 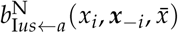, are now obtained by following exactly the same line of argument as in the population genetic model, but using the instrumental distribution instead of the contextual genetic state distribution. Accordingly, we first note that eq. (C.13) says that we predict fitness with the same linear regression

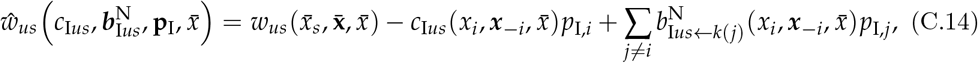

where vector 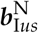 collects the *b*_I*us←k*(*j*)_ regression coefficient and vector **p**_I_ collects all the *p*_I,*i*_ frequencies. Second, we let ***σ**_s_*(*x*_*i*_, ***x**_−i_*, ***p***) and 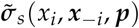 denote the distribution of group states in a population where the vector of frequencies of allele I across classes is equated to the vector ***p*** of mutant allele frequencies across classes in the population genetic model (the ***σ**_s_*(*x*_*i*_, ***x**_−i_*, ***p***) and 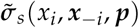 distributions have, respectively, sample space 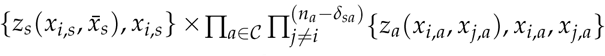 and 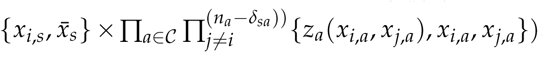. These sample spaces induce the sample simplexes as the 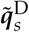 and 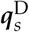 distributions (used in eq. B.16) and so we set

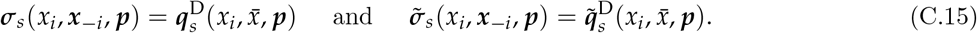

We can now define the quadratic expression

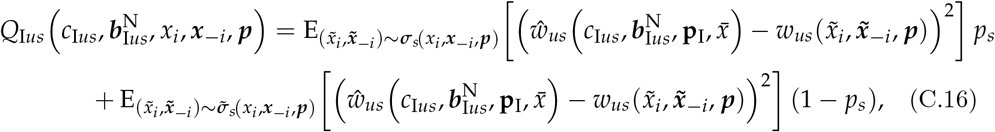

and we solve 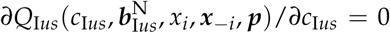 and 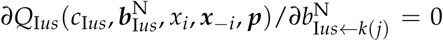 for all *j* ≠ *i*. Recalling eq. (B.20), we obtain the regression coefficients of eq. (C.13) as

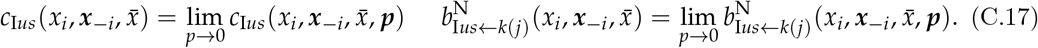

We now further note that when 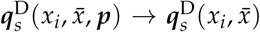 and 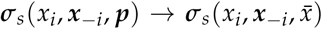 as *p →* 0, we have

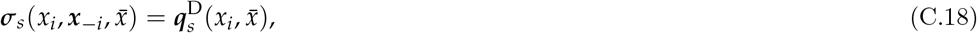

and

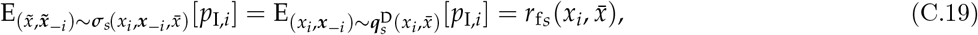

where the first equality follows from eq. (C.18) and the definition of *p*_I,*i*_, which takes exactly the same frequency as the mutant allele within an individual under assumption (C.18); while the second equality follows from the definition of the within-individual identity in state under neutrality (recall eq. B.22). Likewise, when individual *j* ≠ *i* is of class *a*, we have

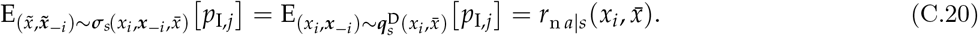

The first equality follows from eq. (C.18) and the definition of *p*_I,*j*_, while the second equality follows from the definition of the between-individual identity in state under neutrality and the fact that the distribution of allele I is the same as that of the mutant allele (recall eq. B.23). In terms of the 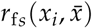 and 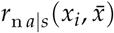 probabilities, we have that

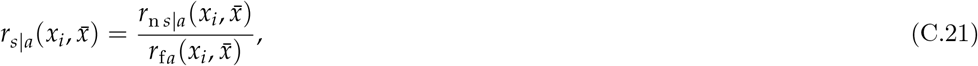

which is the (neutral) relatedness coefficient between a class-*s* actor and a class-*a* recipient in a population monomorphic for trait value 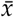.

By contrast to the regression coefficients in the population genetic model, the coefficient
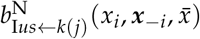 (eq. C.17) takes only the interpretation of a neighbor-modulated regression coefficient, since it gives the effect, on the fitness of *i*, of an average social partner switching to expressing trait *x*_*i*_ instead of its own trait *x*_*j*_. This in general is not the effect of individual *i* on the fitness of its average social partner when switching its own trait value from 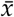 to *x*_*i*_ (since in general 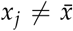). We next show how to obtain an actor-centered regression coefficient out of 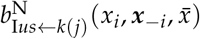.

#### C.3.2 Actor-centered regression coefficients

To introduce our argument, we first consider haploids without class structure. In that case, we set 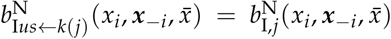 since there are no class effects. It will also turn out useful to re-write the quadratic expression (eq. C.16) for the haploid case in more compact form. To that end, let ***σ***_I_(*x*_*i*_, ***x**_−i_*, *p*) denote the full unconditional instrumental distribution (which concatenates the unconditional instrumental distributions ***σ***(*x*_*i*_, ***x**_−i_*, *p*) and 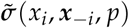 weighted by the respective allele I frequencies), and that has sample space

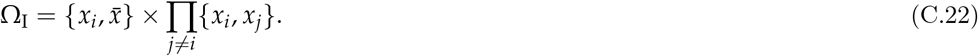

With this, we can then write the quadratic expression for haploids as

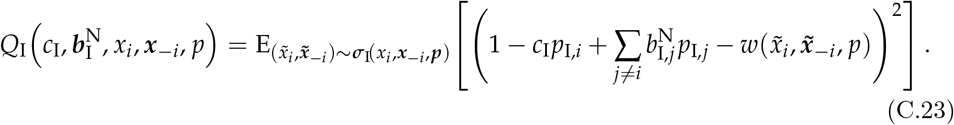

Eq. (C.23) illustrates that, given that we assign a distribution to the traits expressed by different individuals (the ***σ***_I_ distribution), the quadratic expression to be minimized can be represented as an average over a distribution of the traits. The resulting regression coefficients are contrasts of the fitness values (i.e., weighted “averages” except that the sum of the weights is zero), which can no longer be interpreted as averages over the ***σ***_I_ distribution, but whose contrast weights are still fully determined by this distribution (as implied by the so-called“normal equation” of linear regression). For instance, by performing from eq. (C.23) the calculations detailed in the last section, we obtain in the case *n* = 2 (where the sample space is 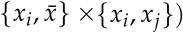,

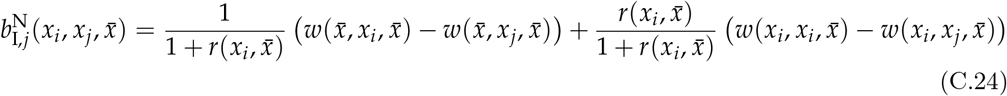

in the limit *p →* 0, which can be thought of as the extension of *b*(*x*, *y*) in eq. (B.11) to the case where the trait of each group member is distinguished.

When individuals are exchangeable (meaning that the distribution of their traits are exchangeable, and that the same fitness function holds for all individuals), then this effect is also an average *b*_I,*j*_ of effects of the focal individual on the *j*th neighbor’s fitness (and thus 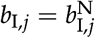). This case holds when one considers mutant-resident dynamics (whether the resident’s state is an uninvadable state, or not) and underlies the dual interpretation of the regression coefficients discussed below eq. (B.7). Otherwise, even if the instrumental distribution ***σ***_I_ is exchangeable between individuals, the traits (sample space) expressed by individuals that do not bear the I allele are not generally exchangeable. For instance, in the above haploid case, the trait sample space of individual *i* is 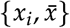 and {*x*_*i*_, *x*_*j*_} and that of its neighbor *j* is {*x*_*i*_, *x*_*j*_}. So these are not exchangeable and we do not have 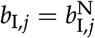.

Rather, in order to construct an additive effect *b*_I,*j*_ of the focal individual on the *j*th neighbor’s fitness, we let *b*_I,*j*_ be the regression coefficient to *p*_I,*j*_ in the quadratic expression eq. (C.23), still under the instrumental distribution defined from the context of individual *i*, but when the traits expressed by the focal and its neighbor *j* are exchanged (in general, this is also different from the regression coefficient to *p*_I,*j*_ under the instrumental distribution defined from the context of individual *j*). For example, consider that *x*_*j*_ is equal to *x*_*i*_, except that we still denote it *x*_*j*_. Then, under the instrumental distribution, *p*_I,*j*_ has no effect on the focal individual’s fitness, since it has no effect on expressed trait, *x*_*i*_ = *x*_*j*_. Yet, the focal individual has a distinct effect on its *j*-neighbor depending on its own *p*_I,*j*_ and this must be reflected by a non-zero *b*_I,*j*_ in the inclusive fitness as-if of individual *i*. This effect is recovered by switching the supports of the focal’s trait and of the *j*-neighbor’s trait, so that the focal trait now has sample space {*x*_*i*_, *x*_*j*_}, but more importantly the neighbor’s trait now has sample space 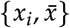 so that we can compute a non-zero *b*_I,*j*_ as the regression coefficient of focal fitness on neighbor’s *p*_I,*j*_.

Formally, this means that once we have an expression, for 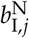 from the regression of *i*’s fitness under the instrumental distribution, as a contrast of individual fitnesses in different contexts, we obtain *b*_I,*j*_ by keeping the contrast weights constant, but modifying the fitness values by switching the sample spaces of the traits expressed by the I allele in individuals *i* and *j*. In the exemplar case *n* = 2 given above (eq. C.24), the switch of the sample spaces 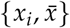 and {*x*_*i*_, *x*_*j*_} then leads to simply interchanging 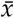 and *x*_*j*_ in eq. (C.24). This produces

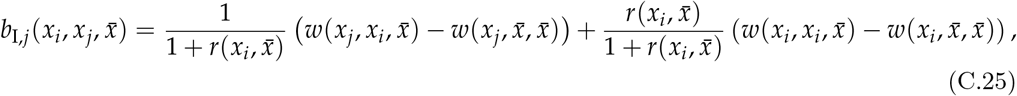

which is an effect of the focal individual on the *j*th neighbor’s fitness. In terms of this coefficient, we can then define the inclusive fitness as-if of individual *i* as

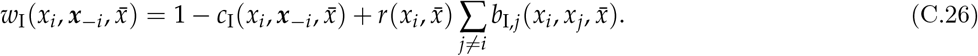

More generally, for diploidy and class-structure the same argument applies. Here, once we have the neighbor-modulated coefficient 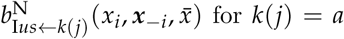 from the regression of *i*’s fitness under the instrumental distribution, as a contrast of individual fitnesses in different contexts, we can obtain 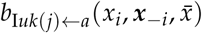 as the effect of an individual *i* of class *a* on receptor *j* of class *k*(*j*) = *s* by keeping the contrast weights constant, but modifying the fitness values by switching the sample spaces of the traits expressed by the focal in class *s* and its neighbor *j* when of class *a*; that is, by switching the supports of the focal’s trait and of the *j*-neighbor’s trait, so that the focal’s trait (now of class *s*) has support 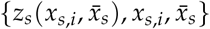 and the neighbor’s trait (now of class *a*) has support {*z*_*a*_(*x*_*i,a*_, *x*_*j,a*_), *x*_*i,a*_, *x_j,a_*}. This then leads a non-zero 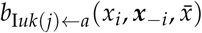 additive effect of a focal of class *a* on its neighbor *j* of class *k*(*j*) = *s*, whose sum over all recipients of class *s*, denoted

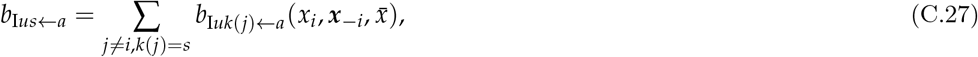

is an individual-centered version of eq. (B.31). In terms of the so-obtained actor-centered indirect coefficient (and the *c*_I*ua*_ and 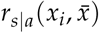) coefficients computed in the previous section, recall eq. C.17 and eq. C.21), we can define the inclusive fitness as-if of an individual *i* of class *a* as

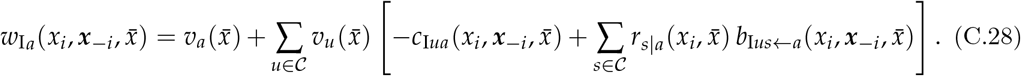

### C.4 Connecting the gene-centered and the rational-actor centered perspectives

Suppose now that at an uninvadable state *x*^∗^, we have

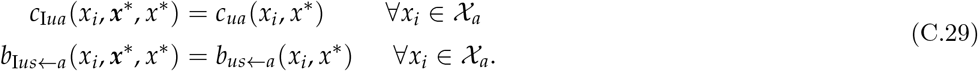

Then, the inclusive fitness as-if at *x*^∗^ is equivalent to the inclusive fitness at *x*^∗^:

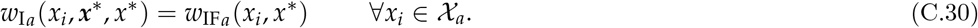

Since we know the right-hand side is maximized at *x*^∗^, i.e., eq. (B.39) is satisfied, it follows from eq. (C.30) that the inclusive fitness as-if (eq. C.28) satisfies eq. (C.3). Thus, in an uninvadable population state, it is as if each individual aimed to maximize its inclusive fitness as-if, when all others exhibit fitness-maximizing behavior.

We now show that eq. (C.29) indeed holds. First, we have both *c*_I*ua*_(*x*_*i*_, ***x***^∗^, *x*^∗^) = *c*_*ua*_(*x*_*i*_, *x*^∗^) and 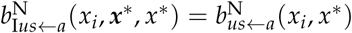 because the regression model eq. (C.13) is the same as that in the population genetic model (eq. B.14), the only difference is that its regression coefficients are evaluated under a distribution where group neighbor traits are all distinct, everything else remaining the same. Hence, in a population where all neighbors of the same class of an individual have the same trait value, the regression coefficients computed under the instrumental and contextual distributions will be the same. It now remains to show that 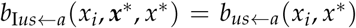. This follows by noting that in the computation leading to 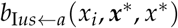, the only thing that is changed, is an exchange of supports in the evaluation of fitnesses, but the position of the argument *x*_*i*_ remains unchanged in all fitness functions. The nature of the maximization problem remains henceforth unchanged, because it is only the trait of neighbors that are interchanged. This has no effect on the symmetric Nash equilibrium (eq. C.3) and so when all neighbors play the same trait, the regression coefficients will be the same. Hence, in an uninvadable population state, it is as-if individuals in each class maximize (in the best-response sense) their own inclusive fitness defined by eq. (C.28).

As explained in the main text, eq. (C.28) is, however, not an operationally convincing as-if fitness as it entails that an individual controls the instrumental distribution describing the number of group neighbors expressing the same trait as self. But since the contextual distribution determining the instrumental distribution (by way of eq. C.18) is a population-level property that depends on the mutant trait, we cannot meaningfully view this distribution as under the control of a particular individual.

### C.5 Weak selection

#### C.5.1 Weak-selection concepts

We now finally turn to deriving an inclusive fitness as-if functions under weak selection (see section “Weak selection concepts” of the main text for an informal discussion). To take weak selection more formally into account, we let the matrix ***A***(*x*, *y*) describing the growth of the mutant when rare in the population with classes (eq. A.8 with elements giving the expected number of groups in state **i** that descend from a group in state **j**) be of the form

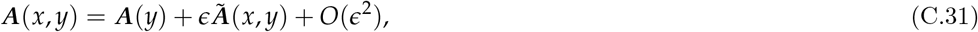

where matrix ***A***(*y*) has leading positive eigenvalue equal to 1 and is independent of the mutant trait, 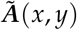 is a matrix depending on both mutant and resident traits, and *e* is a small parameter.

The representation given in eq. (C.31) captures the two kinds of weak selection that we discussed in section “Weak selection concepts” of the main text^3^. First, one can consider that the parameters determining both mutant and resident phenotypic effects are small. In this case of “small-mutation” selection, the matrix ***A***(*y*) depends only the resident trait *y* and matrix 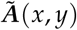 is a first-order polynomial in mutant trait *x*, in which case one can use the approximation ***q***(*x*, *y*) *∼ **q***(*y*). Second, one can consider traits affecting some material payoff (e.g., calory intake), or any other phenotypic feature, which itself affects only weakly a background reproduction and survival (“small-parameter” weak selection). For this case, matrix ***A***(*y*) *→ **A*** is actually independent of both mutant and resident traits, in which case one can use the approximation ***q***(*x*, *y*) *∼ **q*** and ***ϕ***(*x*, *y*) *∼ **ϕ***, and the perturbation matrix 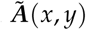 can take any form.

For weak selection, *ϵ →* 0 (e.g, Nagylaki, 1993; Lessard and Soares, 2016), the remainder *O*(*ϵ*^2^) in eq. (C.31) is neglected and

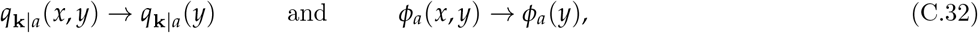

where the left-hand sides depend at most on the resident traits and where **k** can describe either a haploid or diploid group state (in the latter case, **k** must account for heterozygotes and homozygotes within each class), and is independent of the evolving traits altogether under “small-parameter” weak selection. Eq. (C.32) follows from Lessard and Soares (2016, eqs. 59–67) who show that when *ϵ →* 0, the distribution over states of the mutant when rare in the population is described by the right unit eigenvector ***u***(*y*) of ***A***(*y*), and this vector subtends *q*_**k**|*a*_(*y*) and *ϕ_a_*(*y*) (e.g., 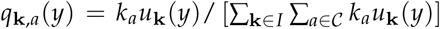, eq. A.21, eq. A.25 and explanations below eq. A.18 for the haploid case). Hence, not only the reproductive value *v*_*u*_(*y*) but also the genealogical and class structure no longer depend on mutant traits. In other words, the population-level properties may vary with the resident trait but are held constant on variation of the mutant trait. This argument also applies to the case of distinct individuals (remember Section A.1.2).

By collecting all components *q*_**k***|a*_(*x*, *y*) into the distribution 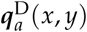 of genetic group states, we have for weak selection that

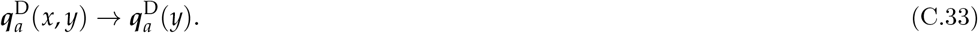

We will next apply eq. (C.33) to derive explicit expressions for fitness as-if under weak selection.

We are now ready to derive explicit as-if fitness representations. Fully endorsing weak selection, we denote from now on by 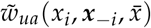 the weak-selection approximation of the class-specific fitness function 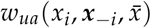 (as this covers both types of small-mutant and of small-parameter weak selection). We first recover two expressions for direct fitness as-if from the literature, which will allow us to point to limitations of this maximand, and finally to formalize inclusive fitness as-if.

#### C.5.2 Class-specific inclusive fitness as-if maximization under weak selection

First, recall that, regardless for any strength of selection, we showed that each individual of each class will appear to maximize its inclusive fitness defined by eq. (C.28) in an uninvadable population state (e.g., eq. C.30, which implies that eq. (C.3) is satisfied) regardless of the the strength of selection. Hence, eq. (C.28) will also be maximized under weak selection (eq. (C.3) is satisfied), with the only difference that we can let the population genetic structure be independent of the actor’s traits. Hence, for weak selection we write the inclusive fitness as-if of individual *i* of class *a* as

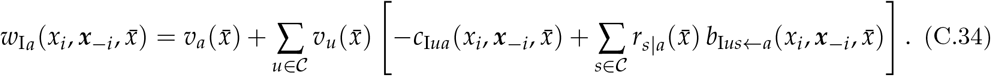

This is eq. (6) of the main text and the key difference with eq. (C.28) is that relatedness is now independent of the actor’s trait and the regression coefficients are evaluated under weak selection using the fitness function 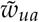 (instead of the fitness function *w*_*ua*_) and using the weak selection approximation of the instrumental distribution, everything else remains the same.

## Footnotes

1 Strictly speaking, Grafen (2006a) did not formally prove any form of inclusive fitness maximization, since as discussed in Lehmann and Rousset (2014a), he considered selection on a mutant allele over a single generation under the assumption that there is a single copy of this allele in the population. But his results about individual-centered inclusive fitness maximization can be shown to hold by considering multigenerational effects and are implied by the present analysis.

2 We assume that is a locally convex Hausdorff space; namely, it is a nonempty, compact, and convex set in a topological vector space (Alipantris and Border, 2006, p. 55). We are not aware of any applications in evolutionary biology that is not covered by this case (e.g., it covers discrete finite trait sets, infinite-dimensional reaction norms or function-valued traits, combination of thereof, etc.), and is the space for which general results concerning function maximization exists (Alipantris and Border, 2006, pp. 581-585).

3 As concrete example of both “small-mutation” and “small-parameter” weak selection, we can use the social-insects scenario and corresponding fitnesses given in Box 1. Then, we can first Taylor-expand the fitness components, say the number of daughters produced by queens (eq. B.2), in mutant trait around the resident trait and neglect higher-order terms to obtain 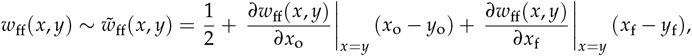 where the right-hand side gives a small-mutation approximation to fitness as the fitness of a resident individual in a monomorphic resident population plus the marginal changes in fitness weighted by their phenotypic differences. Alternatively, we can linearize fitness in terms of the effect *P*(*x*_o_) of workers on female fecundity 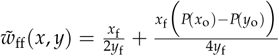 where the right-hand side represents small-parameter approximation to fitness.

